# An optimised approach to evaluate variability in gut health markers in healthy adults

**DOI:** 10.1101/2024.07.25.604267

**Authors:** Kirsten Krüger, Yoou Myeonghyun, Nicky van der Wielen, Dieuwertje Kok, Guido J. Hooiveld, Shohreh Keshtkar, Marlies Diepeveen-de Bruin, Michiel G.J. Balvers, Mechteld Grootte-Bromhaar, Karin Mudde, Nhien T.H.N Ly, Yannick Vermeiren, Lisette C.P.G.M de Groot, Ric C.H. de Vos, Gerard Bryan Gonzales, Wilma T. Steegenga, Mara P.H. van Trijp

## Abstract

Despite advances in gut health research, the variability of important gut markers within individuals over time remains underexplored. We investigated the intra-individual variation of various faecal gut health markers using an optimised processing protocol aimed at reducing variability. Faecal samples from ten healthy adults over three consecutive days demonstrated marker-specific intra-individual coefficients of variation (CV%), namely: stool consistency (16.5%), water content (5.7%), pH (3.9%), total SCFAs (17.2%), total BCFAs (27.4%), total bacteria and fungi copies (40.6% and 66.7%), calprotectin and myeloperoxidase (63.8% and 106.5%), and untargeted metabolites (on average 40%). For thirteen microbiota genera, including *Bifidobacterium* and *Akkermansia*, variability exceeded 30%, whereas microbiota diversity was less variable (Phylogenetic Diversity 3.3%, Inverse Simpson 17.2%). Mill-homogenisation of frozen faeces significantly reduced the replicates CV% for total SCFAs (20.4% to 7.5%) and total BCFAs (15.9% to 7.8%), and untargeted metabolites compared to only faecal hammering, without altering mean concentrations. Our results show the potential need for repeated sampling to accurately represent specific gut health markers. We also demonstrated the effectiveness of optimised preprocessing of stool samples in reducing overall analytical variability.

## Introduction

Nowadays the impact of gut health on the maintenance of overall human health is commonly acknowledged [1, 2]. So far, gut health research has strongly focused on faecal microbiota composition, because microbiota shifts have been correlated to many diseases [3, 4]. Numerous factors like demographics [5, 6], diet [7–9], physical activity [10, 11], medication use [12, 13], age [14, 15], health status [16], gut transit time [17, 18], and other environmental factors [19] have been reported to affect the microbiota. Interestingly, only a minor fraction of inter-individual microbiota differences can be explained by such exogenous and intrinsic host factors [20].

To provide a comprehensive characterisation of an individual’s gut health status, it is of interest to measure a combination of gut health-related markers. For instance, the Bristol Stool Scale (BSS) can be used as proxy for faecal water content % (WC%) and intestinal transit time (ITT) [17, 21, 22]. Water content is however rarely measured [23, 24]. In addition to relative microbiota abundance, absolute microbiota numbers have also been suggested to be of importance for the interpretation of microbiota data [25–27]. Moreover, important other kingdoms than bacteria, for instance fungi have been described [28, 29]. Until recently, the importance of the intestinal fungal microbiome (“mycobiome”), has been largely unstudied and only constitute 3% of the total microbiome studies [30]. Importantly, while many studies focus on microbiota composition, the microbiota-derived metabolites might be the crucial functional regulators in health and disease [31–35]. One well-known example is the suggested health benefits of short-chain fatty acids (SCFAs) [36]. Furthermore, branched-chain fatty acids (BCFAs) are mainly produced by the microbial fermentation of branched-chain amino acids [37], and receive attention in human studies more frequently, as potential negative health associations have come to light [38, 39]. Faecal pH is an indicator of the intestinal environment, which could be of relevance given its proposed influence on bacterial metabolic processes, growth and composition [39, 40], and is associated with the presence of faecal SCFAs and BCFAs [41, 42]. Despite being easily measured with a probe, pH is infrequently assessed in human studies in the context of gut health. Furthermore, faecal biomarkers that are (potentially) clinically relevant include the inflammatory markers calprotectin and myeloperoxidase (MPO) [43, 44]. Both inflammatory markers are neutrophil proteins, with calprotectin commonly used as a biomarker in inflammatory bowel disease (IBD) [45], while MPO has been highlighted in recent years, as a potential biomarker for inflammation [46].

As yet, the intra- and inter-individual variation of most of the abovementioned gut health markers have not been well studied in literature. Data from healthy subjects [47, 48] and patients [49] highlight the importance of repeated sampling, with three to five consecutive faecal samplings, to capture the intra-individual microbiota variation. Additionally, large day-to-day variations in the microbiota composition and diversity within individuals in consecutive faecal samples over a six-week period were found [48]. A longitudinal study suggested that faecal microbiota is generally stable over one year, but can undergo rapid and profound changes due to factors like traveling [50]. Even in controlled enclosed environments, the human faecal microbiota was found to be dynamic [51].

Additionally, for faecal metabolite evaluation it might be relevant to include repeated faecal samples, because the three main SCFAs acetate, propionate, and butyrate showed considerable intra-individual variability in four human faecal samples collected within a week [52]. Also, the total SCFAs, and especially butyrate, varied over 12 weeks in free-living adults [53]. Moreover, stool consistency variability is high in both inflammatory bowel syndrome patients and healthy subjects [49]. The variability of other gut health markers in consecutive collected faecal samples is less well-described. More in-depth knowledge about intra-individual variation in gut health markers enhance understanding of the relevance of repeated faecal measurements to control for temporal variation. Furthermore, establishing more accurate baseline values of these markers within participants is crucial in assessing the actual intervention-induced effects, rather than studying day-to-day variation.

Limitations and differences in faecal sampling- and handling methodology may play a role in the large inter-individual differences that are reported [54, 55]. Currently, there are no clearly defined standard practices for faecal sampling, handling, and analysis in human studies. Often only a single scoop of a few grams of faeces is collected [56]. Optimising gut health analysis using faeces might require adjusted sampling and processing procedures. For example, many microbiota-derived metabolites are volatile, sensitive to degradation, and require specific solvents [54, 55, 57–63]. To measure the microbiota, microbial metabolites, and other gut markers accurately and with minimal technical variation, strict faecal sample collection, storage, preparation, and analytical procedures are required [54, 64]. These procedures are especially important to reduce variation due to sample handling, which could be falsely attributed to biological intra-and inter-individual differences, or give a false indication of the baseline versus post-intervention effects on gut health status [56].

One important consideration in the sampling procedures is the collection of larger volumes of faeces by the participant by taking multiple scoops from different locations of the faeces, because it was previously shown that spot sampling rather than collection from the top, middle, bottom or edge positions of the faeces resulted in higher microbiota and metabolites variability [60, 65]. Other examples of optimised procedures include keeping samples frozen during processing at all times, thus avoiding freeze-thaw cycles and temperature fluctuations which may lead to metabolite degradation and microbial fermentation, and selecting appropriate storage conditions [54, 66]. Furthermore, because of the heterogeneity of faeces [60, 65], applying specific devices to properly homogenise the samples becomes crucial. Blenders [64] or other milling devices suitable of grinding deep-frozen materials into fine powders, such as an IKA mill, are often used in plant metabolomics and soil microbiome research [67], while these are not often reported in the context of human faeces processing. Nevertheless, homogenising faeces may reduce the variation in bacteria abundances and SCFAs levels as compared to non-homogenised faeces [65, 68].

We investigated the variation of a selected panel of gut markers in healthy adults, with a focus on the intra-individual variation. Furthermore, the effect of an optimised sample pre-processing procedure, including mill-homogenisation in liquid nitrogen, on the analysis of these gut health markers was evaluated. For this purpose, ten participants collected three faecal samples on consecutive days. The optimised faecal sampling, processing and storage protocol was applied. The measured gut health markers included BSS, pH, and WC%, microbiota composition and diversity, absolute abundance of the total bacteria and fungi, SCFAs, untargeted metabolite profiles and specific inflammatory biomarkers.

## Results

### Subjects and faecal sample collection

Ten healthy adults finished the study, of which 2 males and 8 females. Seven participants collected their stools on three consecutive days (**Supplementary Table 1**), while three participants collected on non-consecutive days.

### Variation of a selected panel of gut health markers

The results presented in **Figure 1** reveal the intra-subject variation for a panel of different gut health markers. The BSS and pH demonstrated a low CV%_intra_ of 16.5±14.9 and 3.87±1.74, with a moderate test-retest reliability of ICC 0.74 [0.43-0.92] and ICC 0.56 [0.16-0.85] (**Table 1**). Water content demonstrated a low CV%_intra_ of 5.74±3.20 with a low test-retest reliability of ICC 0.37 [-0.01-0.76]. Faecal SCFAs analysis revealed CV%_intra_ of below 30% for total SCFAs (17.2±13.8), total BCFAs (27.4±15.2), acetic acid (16.0±11.7), propionic acid (17.8±12.4) and butyric acid (27.8±17.4). Isobutyric acid, valeric acid, and 4-methyl valeric acid had a CV%_intra_ lower than 30%, while isovaleric acid, heptanoic acid, and hexanoic acid had a CV%_intra_ higher than 30% (**Table 1**). Test-retest reliability scores showed moderate test-retest reliability for total SCFAs (ICC 0.65 [0.29-0.89]), acetic acid (ICC 0.73 [0.41-0.92]) and propionic acid (ICC 0.64 [0.28-0.88]) while butyric acid (ICC 0.40 [-0.01-0.77]) and total BCFAs (ICC 0.35 [-0.03-0.74]) showed poor test-retest reliability. To assess whether the genetic potential for fermentation and production of SCFAs was less variable compared to the metabolites, microbial pathways and enzymes related to fermentation and production of SCFAs in the predicted bacterial genomes were analysed (**Supplementary Table 2**). The CV%_intra_ for pathways and enzymes related to acetic acid, propionic acid, and butyric acid production were 22.4±22.3, 23.4±13.3, and 23.1±9.43, respectively. From the 26 pathways and enzymes, 14 demonstrated good or excellent test-retest reliability over time (ICCs 0.75-1.0), 10 demonstrated moderate test-retest reliability over time (ICCs 0.50-0.75), and two demonstrated poor test-retest reliability. Furthermore, high CV%_intra_ were demonstrated by both total fungi (67.0±35.8, corrected by wet weight) and total bacteria (40.6±16.9, corrected by wet weight). Total fungi counts showed excellent test-retest reliability (ICC 0.90 [0.75-0.97], corrected by wet weight), while total bacteria showed poor test-retest reliability ICC 0.32 [-0.01-0.71] (corrected by wet weight). Corrections for faecal dry weight demonstrated similar results for both the CV%_intra_ and ICC of the total fungi and total bacteria copies compared to corrections for faecal wet weight (**Supplementary Table 3**). Calprotectin showed a high CV%_intra_ of 63.8±69.0 with excellent test-retest reliability (ICC 0.92 [0.79-0.98]), and MPO demonstrated a high CV%_intra_ of 106.5±51.1 with poor test-retest reliability (ICC 0.37 [-0.03-0.77]) largely driven by variation in participant 10. All markers demonstrated an intra and inter-assay technical variation of <14% (**Supplementary Table 4**).

**Figure 1.**
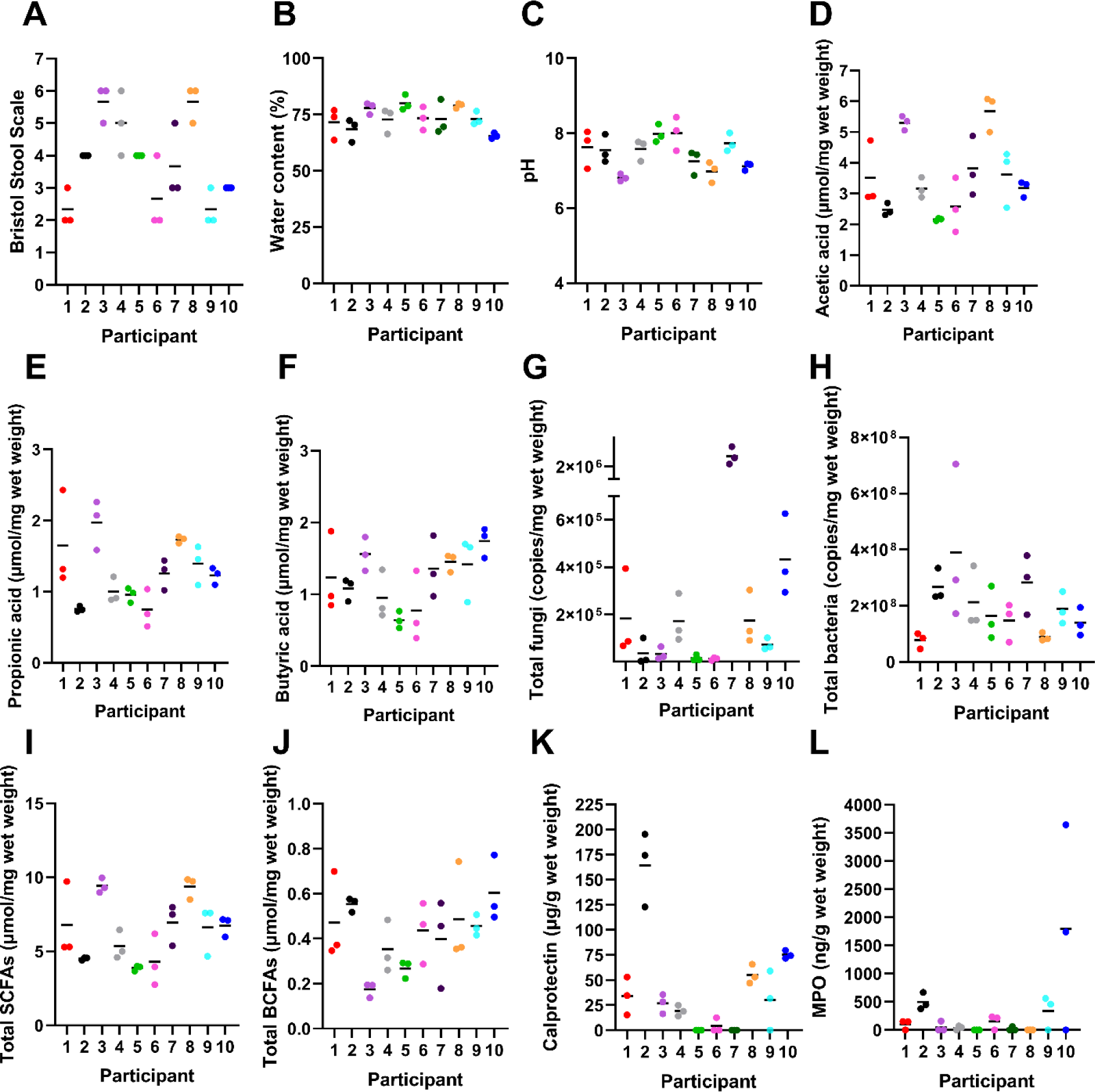
Intra-individual variation in a panel of gut health markers in faecal samples of healthy subjects collected over three consecutive days. The A) Bristol Stool Scale, B) water content, C) pH, D) acetic acid, E) propionic acid, F) butyric acid, G) total fungi, H) total bacteria, I) total SCFAs, J) total BCFAs, K) calprotectin, and L) MPO are presented as measured in the three faecal samples collected. Data from n=10 participants are presented. For calprotectin, n=8 subjects had at least one measurement above the detection limit of 11.92 µg/g wet weight, and for MPO, also n=8 subjects had at least one measurement above the detection limit of 68.5 ng/g wet weight. Individual participant’s samples are shown by the three dots. Participants were instructed to choose only one scale point for the Bristol Stool Scale, but when two types were indicated by the participant, the highest of the two scores was included in the analysis.

**Table 1.** Variability of a panel of gut health markers in faecal samples of healthy subjects collected over three consecutive days.

| Marker | Coefficient of variation % <sup>1</sup> |  | Intra-class correlation coefficient [CI]<sup>2</sup> |
| --- | --- | --- | --- |
|  | Intra-individual (CV%<sub>intra</sub>) | Inter-individual (CV%<sub>inter</sub>) |  |
| BSS | 16.5 ±14.9 | 36.9±1.85 | 0.74 [0.43-0.92] |
| pH | 3.87 ±1.74 | 6.47±1.46 | 0.56 [0.16-0.85] |
| Water content | 5.74 ±3.20 | 8.13±0.64 | 0.37 [-0.01-0.76] |
| Acetic acid | 16.0±11.7 | 36.1±3.10 | 0.73 [0.41-0.92] |
| Propionic acid | 17.8±12.4 | 36.5±8.85 | 0.64 [0.28-0.88] |
| Butyric acid | 27.8±17.4 | 37.9±0.72 | 0.40 [-0.01-0.77] |
| Valeric acid | 27.6±17.5 | 38.6±3.37 | 0.33 [-0.05-0.73] |
| Heptanoic acid | 81.5±65.4 | 108.9±7.76 | 0.73 [0.41-0.92] |
| Hexanoic acid | 38.2± 31.9 | 93.1±6.59 | 0.81 [0.55-0.94] |
| Isobutyric acid | 24.9±14.5 | 36.5±5.65 | 0.34 [-0.05-0.72] |
| Isovaleric acid | 30.5± 16.7 | 44.1±9.78 | 0.35 [-0.03-0.74] |
| 4-Methylvaleric acid | 26.0 ±41.8 | 35.1±0.00 | 0.12 [-0.39-0.37] |
| Total SCFAs<sup>3</sup> | 17.2±13.8 | 34.3±3.39 | 0.65 [0.29-0.89] |
| Total BCFAs<sup>3</sup> | 27.4±15.2 | 40.5±7.97 | 0.35 [-0.03-0.74] |
| Total fungi<sup>4</sup> | 67.0±35.8 | 222.0±16.7 | 0.90 [0.75-0.97] |
| Total bacteria<sup>4</sup> | 40.6±16.9 | 55.0±18.9 | 0.32 [-0.01-0.71] |
| Calprotectin | 63.8±69.0 | 125.8±0.68 | 0.92 [0.79-0.98] |
| Myeloperoxidase | 106.5±51.1 | 158.7±18.0 | 0.37 [-0.03-0.77] |

<sup>1</sup> The mean ±SD of CV% the n=10 participants is shown. <sup>2</sup> The 95% confidence interval is provided.
Participants were instructed to choose only one scale point for the Bristol Stool Scale, but when two types were indicated by the participant, the highest of the two scores was included in the analysis. <sup>3</sup> The total SCFAs include acetic acid, propionic acid, butyric acid, valeric acid, heptanoic acid, hexanoic acid. The total BCFAs include isobutyrate, isovalerate and 4-methylvaleric acid. <sup>4</sup> The total fungi and bacteria are corrected for faecal wet weight.

The CV%_intra_ of faecal water content demonstrated significant positive correlations with the CV%_intra_ of butyric acid and calprotectin (**Supplementary Figure 1**). The CV%_intra_ of stool consistency, assessed by the BSS, demonstrated significant positive correlations with the CV%_intra_ of acetic acid, propionic acid, butyric acid, hexanoic acid, and the total SCFAs. In summary, all markers except for total fungi, total bacteria, calprotectin and MPO, demonstrated a CV%_intra_ of less than 30%. Overall, 8 out of 12 markers demonstrated moderate to excellent test-retest reliability, with poor test-retest reliability shown by water content, butyrate, MPO, total BCFAs, and total bacteria counts (both wet and dry weight corrections).

### Variation in faecal microbiota composition and diversity

The relative microbiota composition was plotted to visualise changes of the top 25 genera at the three consecutive timepoints (**Figure 2**). Visualisation of the overall microbiota community revealed clusters based on individuals and not per sampling timepoint (**Figure 3A**). From the top 25 genera, 13 genera demonstrated CV%_intra_ >30%, with CV%_intra_ ranging between 15.1 and 84.4. For all bacteria that were measured on genus level, the mean CV%_intra_ was 56.5±9.0, of which 43.1±14.7% of the genera had a CV%_intra_ lower than 30% (**Supplementary Table 5**). Furthermore, 17 bacteria were demonstrating good or excellent test-retest reliability over time (ICCs 0.75-1.0), 7 genera demonstrating moderate test-retest reliability over time (ICCs 0.50-0.75), and one genus, *Akkermansia*, demonstrating poor test-retest reliability (ICC 0.48 [0.12-0.81], **Table 2**). Also the faecal microbiota alpha-diversity was quantified (**Figure 2B**), as calculated by the indexes Fisher Diversity, Inverse Simpson, Shannon Diversity, Phylogenetic Diversity, and the estimated number of species (Chao1) that had a CV%_intra_ of 13.9±8.19, 17.2±10.1, 3.42±1.68, 3.27±2.77, and 11.4±5.06, respectively. All demonstrated moderate to good test-retest reliability over time, namely ICC 0.55 [0.17-0.84], 0.77 [0.48-0.93], 0.81 [0.56-0.95], 0.87 [0.68-0.96], and 0.61 [0.25-0.87], respectively. Overall, the microbiota dissimilarity between individuals is much larger compared to the day-to-day microbiota variation within individual. The microbiota alpha-diversity per individual within one week of consecutive sampling remained stable, but the variability of the bacteria was highly genus-dependent.

**Figure 2.**
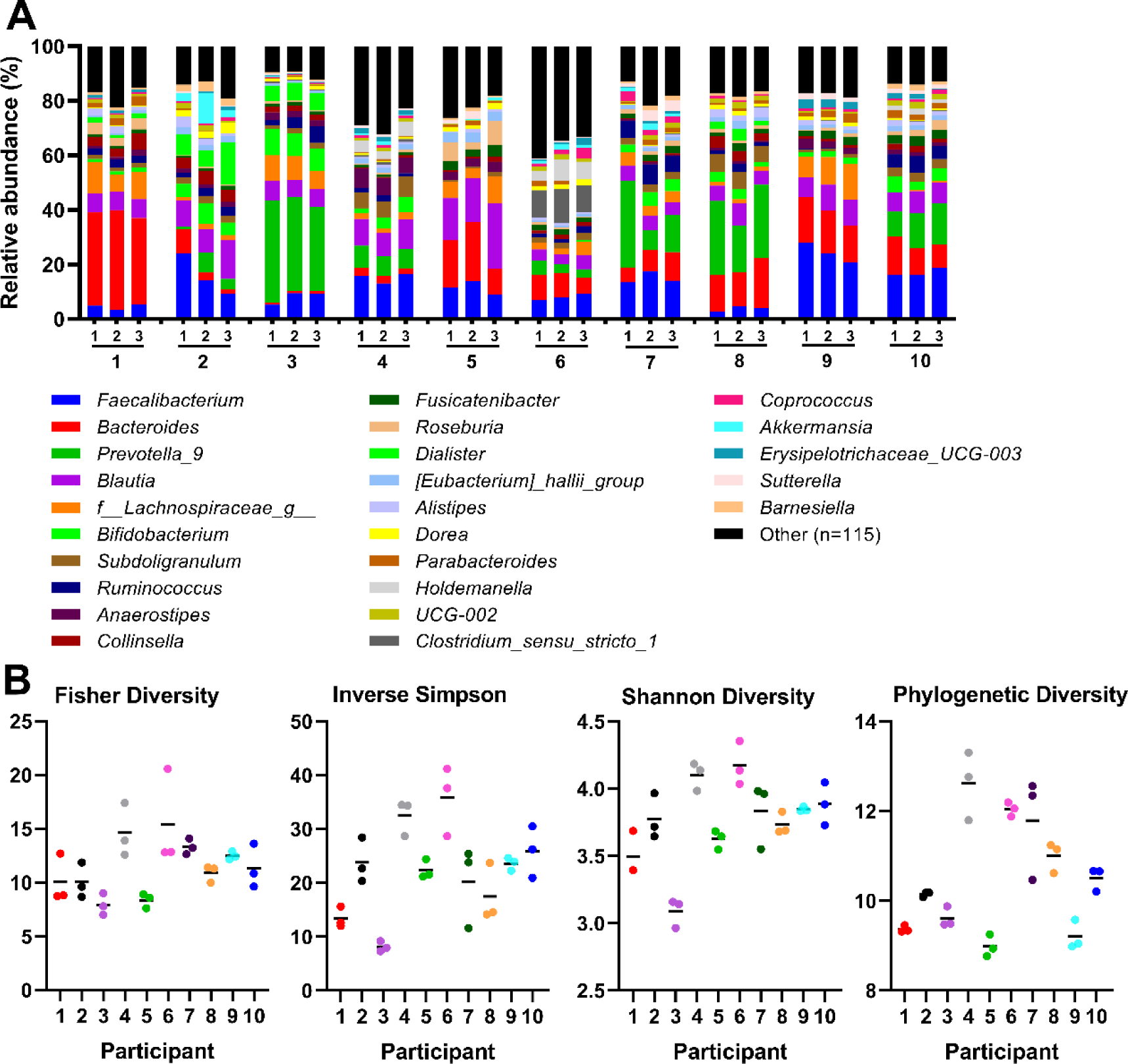
The variation in microbiota composition and diversity in faecal samples of healthy subjects collected over three consecutive days. A) The relative abundance of the faecal microbiota at three consecutive sampling timepoints in the ten individuals. The top 25 bacteria on genus level are shown. If genus was not classified, the family name (f_) is provided. B) The microbiota alpha-diversity, measured by the indexes Fisher Diversity, Inverse Simpson, Shannon Diversity and the Phylogenetic diversity. Individual participant’s samples are shown by the three dots. Data from n=10 participants are presented.

**Figure 3.**
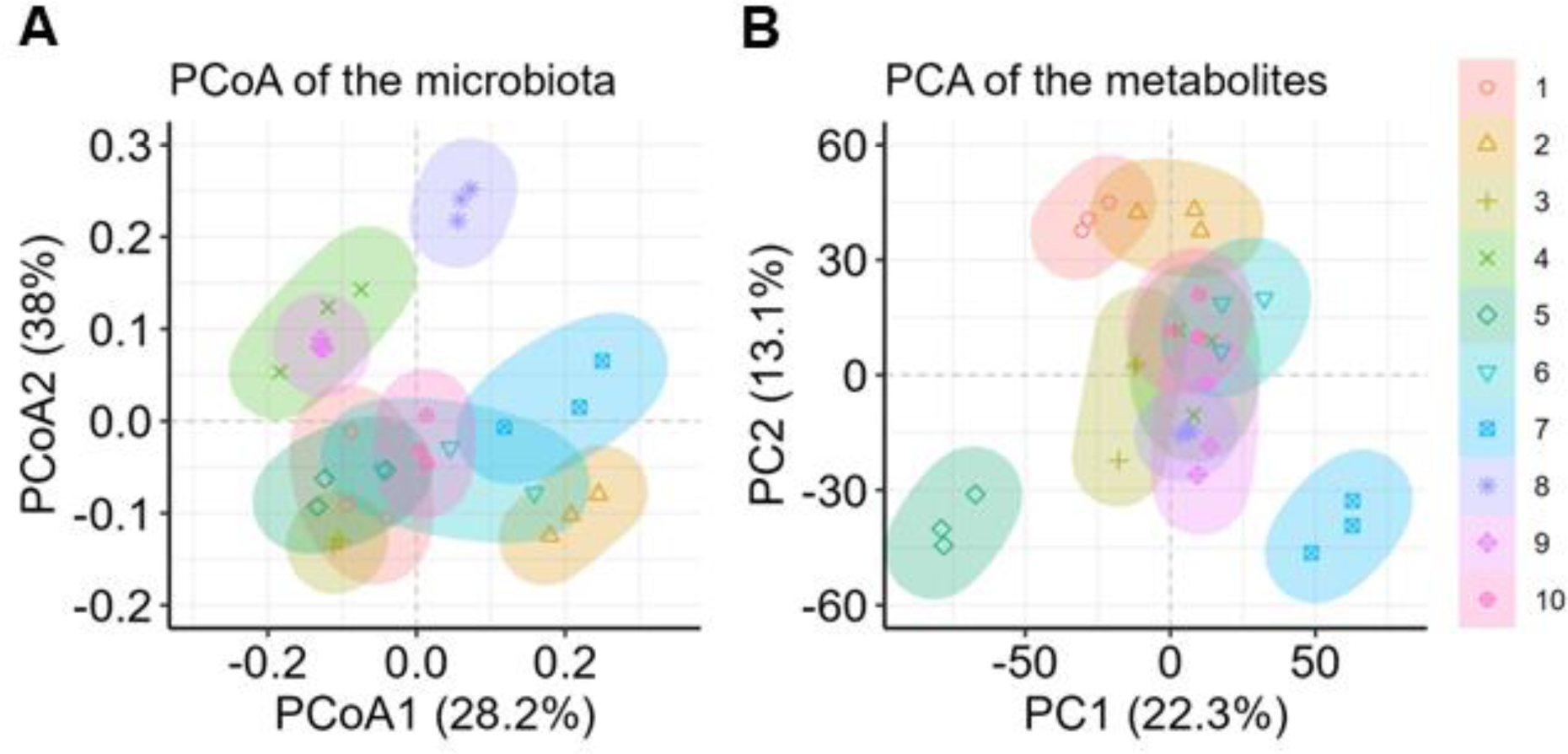
Variation in faecal microbiota and metabolite profiles in faecal samples of healthy subjects collected over three consecutive days. (A) The PCoA plot visualises the microbiota variation between the three consecutive timepoints per subject the beta-diversity was calculated using Weighted UniFrac. (B) The PCA scores plot based on all metabolites measured in both positive and negative ionization modes. The 95% confidence ellipses are shown, coloured according to the ten individuals. PCA: Principal Component Analysis, PCoA: Principal Coordinate Analysis.

**Table 2.** The variation in the top 25 bacteria on genus level and microbiota diversity in faecal samples of healthy subjects collected over three consecutive days.

|  | Coefficient of variation <sup>1</sup> % |  | Intra-class correlation coefficient [CI]<sup>2</sup> |
| --- | --- | --- | --- |
|  | Intra-individual (CV%<sub>intra</sub>) | Inter-individual (CV%<sub>inter</sub>) |  |
| Bacteria<sup>3</sup> |  |  |  |
| *Faecalibacterium* | 21.5±11.4 | 54.7 ± 9.33 | 0.79 [0.52-0.94] |
| *Bacteroides* | 28.1±23.4 | 86.0 ± 7.48 | 0.91 [0.76-0.97] |
| *Prevotella_9* | 35.7±29.8 | 117.7 ± 4.29 | 0.85 [0.64-0.96] |
| *Blautia* | 15.1±9.21 | 48.4 ± 12.9 | 0.79 [0.53-0.94] |
| *f_Lachnospiraceae_g* | 29.6±18.9 | 79.1 ± 3.14 | 0.80 [0.54-0.94] |
| *Bifidobacterium* | 45.1±49.9 | 84.9 ± 9.18 | 0.81 [0.60-0.95] |
| *Subdoligranulum* | 15.3±21.0 | 69.5 ± 4.93 | 0.96 [0.89-0.99] |
| *Ruminococcus* | 26.5±17.3 | 82.8 ± 7.03 | 0.84 [0.61-0.95] |
| *Anaerostipes* | 25.3±17.6 | 88.4 ± 5.53 | 0.93 [0.81-0.98] |
| *Collinsella* | 29.1±23.1 | 80.1 ± 0.58 | 0.85 [0.63-0.96] |
| *Fusicatenibacter* | 21.0±10.8 | 48.8 ± 2.52 | 0.77 [0.49-0.93] |
| *Roseburia* | 36.7±14.6 | 94.9 ± 25.6 | 0.69 [0.37-0.90] |
| *Dialister* | 35.2±19.5 | 168.2 ± 24.0 | 0.79 [0.52-0.94] |
| *[Eubacterium]_hallii_group* | 15.4±12.7 | 55.3 ± 5.43 | 0.90 [0.73-0.97] |
| *Alistipes* | 41.2±27.6 | 72.6 ± 14.5 | 0.60 [0.23-0.87] |
| *Clostridium_sensu_stricto_1* | 84.4±62.1 | 264.7 ± 7.82 | 0.98 [0.95-1.00] |
| *Dorea* | 15.5±6.71 | 64.5 ± 8.77 | 0.86 [0.65-0.96] |
| *Parabacteroides* | 38.5±25.5 | 73.9 ± 13.8 | 0.65 [0.29-0.89] |
| *Holdemanella* | 31.3±3.72 | 186.4 ± 11.7 | 0.88 [0.69-0.96] |
| *UCG-002* | 32.5±17.1 | 61.8 ± 2.24 | 0.69 [0.37-0.90] |
| *Coprococcus* | 29.2±19.4 | 84.3 ± 15.7 | 0.59 [0.23-0.86] |
| *Akkermansia* | 82.0±45.0 | 161.8 ± 29.2 | 0.48 [0.12-0.81] |
| *Erysipelotrichaceae_UCG-003* | 45.8±38.0 | 98.5 ± 12.4 | 0.69 [0.34-0.90] |
| *Sutterella* | 36.7±26.4 | 106.6 ± 27.8 | 0.67 [0.32-0.89] |
| *Barnesiella* | 49.3±51.1 | 101.8 ± 5.16 | 0.85 [0.60-0.96] |
| Alpha-diversity and richness |  |  |  |
| Fisher Diversity | 13.9±8.19 | 25.1 ± 6.31 | 0.55 [0.17-0.84] |
| Inverse Simpson | 17.2±10.1 | 40.2 ± 8.33 | 0.76 [0.48-0.93] |
| Shannon Diversity | 3.42±1.68 | 8.81 ± 0.69 | 0.81 [0.56-0.94] |
| Phylogenetic Diversity | 3.27±2.77 | 12.8 ± 1.09 | 0.87 [0.68-0.96] |
| Chao1 | 11.4±5.06 | 20.1 ± 1.68 | 0.61 [0.25-0.87] |

<sup>1</sup> The mean ±SD of the n=10 participants is shown. <sup>2</sup> The 95% confidence interval is provided. <sup>3</sup> If genus was not classified, the family name (f\_) is provided.

### Variation of untargeted metabolites

In addition to targeted metabolite analyses, untargeted metabolomics was also applied to investigate the variation of all features, which are signals representing chemical compounds measured in the positive or negative ionization mode, in the faecal samples. The number of detected features, the pre-processing workflow, and final included features can be found in **Supplementary Figure 2**. As shown in **Figure 3B**, the differences in metabolites within individuals are smaller than differences between individuals. For the positive features, the mean CV%_intra_ was 40.1±4.44, of which 49.1±6.8% of the features had a CV%_intra_ lower than 30% (**Table 3**). 15.2% demonstrated good or excellent test-retest reliability over time (ICCs 0.75-1.0), 37.0% demonstrated moderate test-retest reliability over time (ICCs 0.50-0.75), and 47.8% demonstrated poor test-retest reliability (ICC<0.50, **Table 3**). For the negative features, the mean CV%_intra_ was 41.7±5.43, of which 47.6±7.6% of the features had a CV%_intra_ lower than 30%. In comparison, the CV%_inter_ was higher than the CV%_intra_, namely 76.9±2.18 for the positive features, and 89.0±3.52 for the negative features. 24.1% demonstrated good or excellent test-retest reliability over time (ICCs 0.75-1.0), 37.3% demonstrated moderate test-retest reliability over time (ICCs 0.50-0.75), and 38.7% demonstrated poor test-retest reliability (ICC<0.50).

**Table 3.**
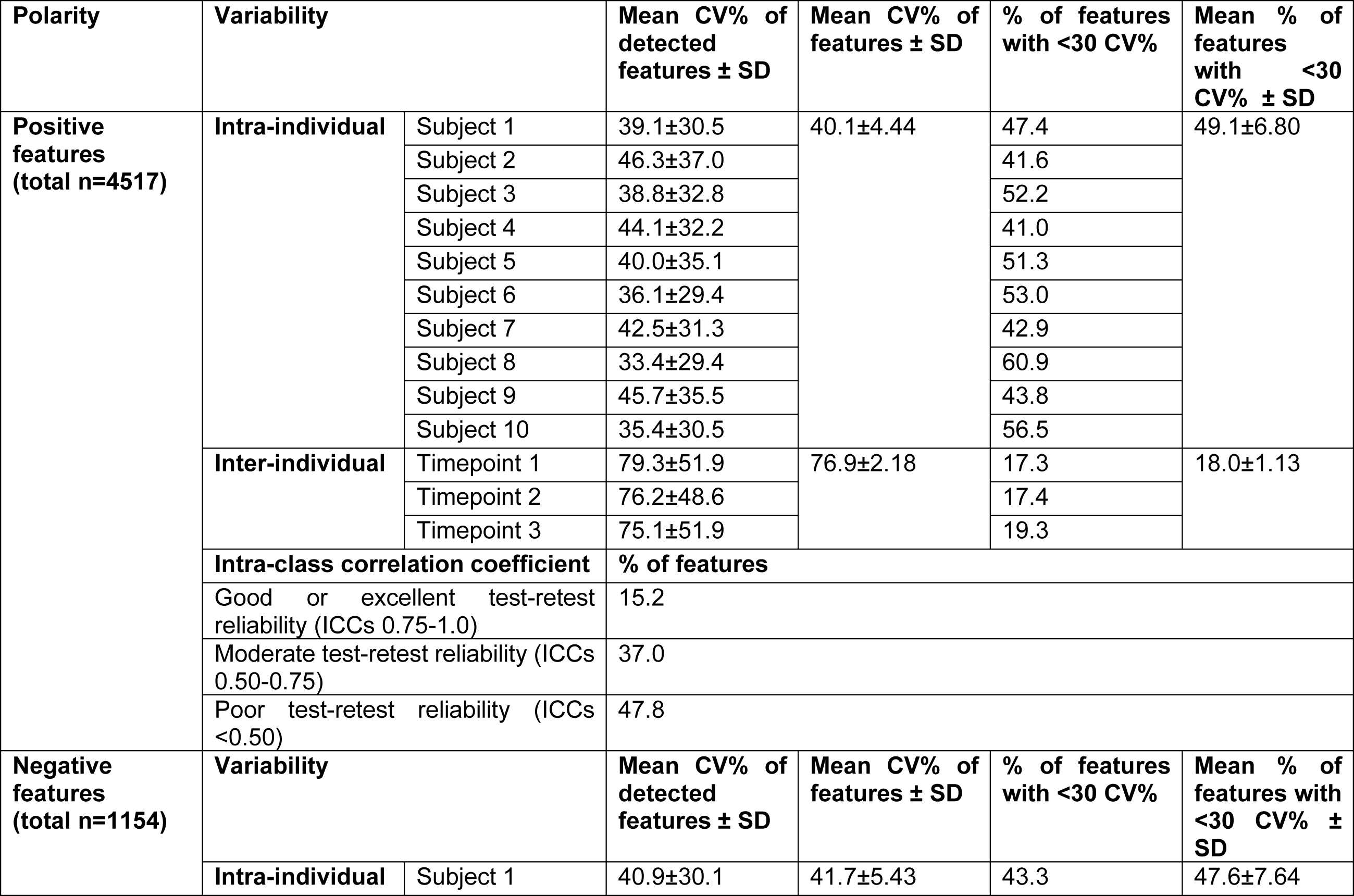
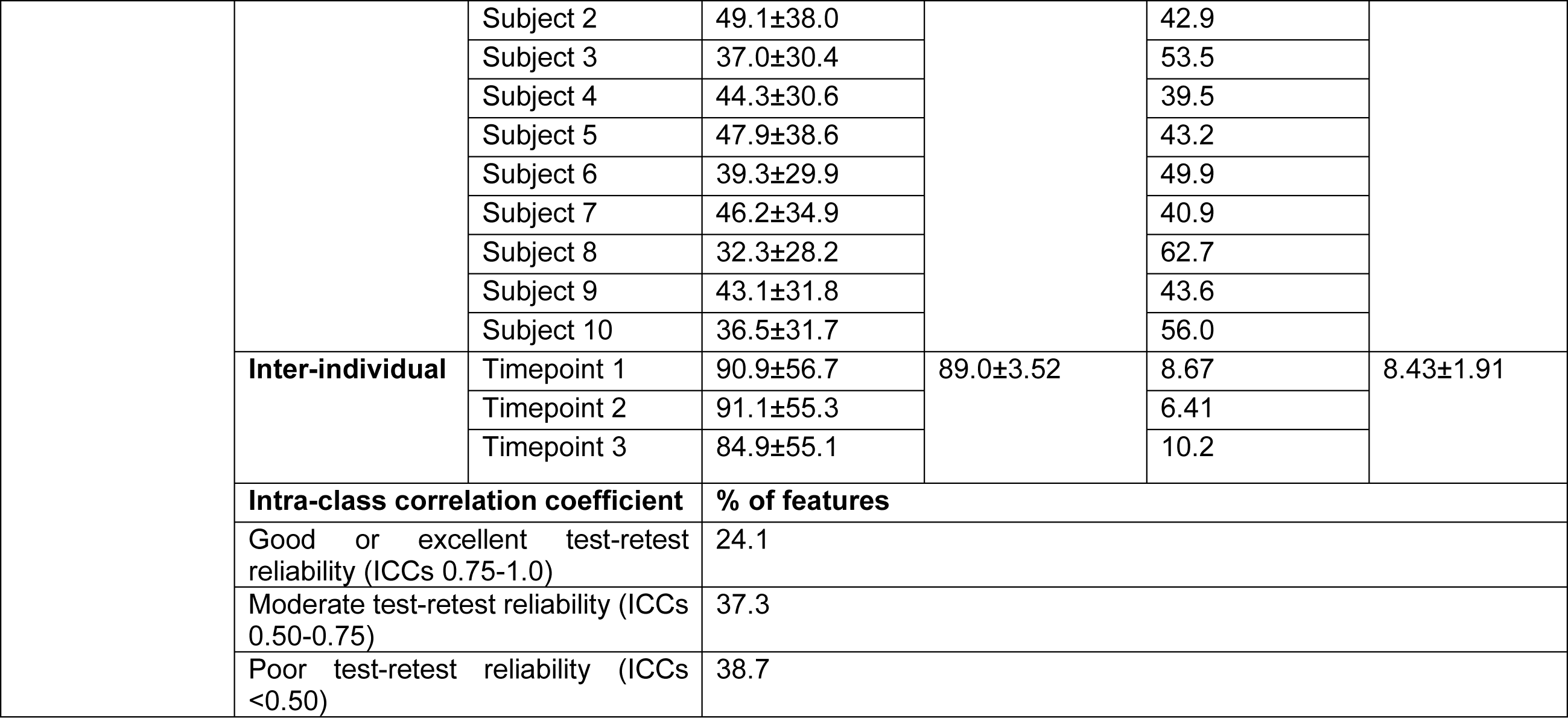
The variability of the metabolites in faecal samples of healthy subjects collected over three consecutive days expressed as measured with coefficient of variation and the inter-class correlation coefficient.

### The effect of faecal sample preparation on the gut-health related outcome measures

To determine the most appropriate pre-processing procedure per marker, the effect of faecal sample homogenisation on the analysis of gut-health markers was investigated using ten replicates from the same pool of faeces. Five of these replicates were processed by hammering only, while the other five replicates were mill-homogenised after hammered (**Figure 4A**). Both pH and water content showed a slight reduction in CV% due to the additional mill homogenisation: for pH from 0.43% (hammered-only) to 0.38% (mill-homogenised), and for water content from 1.63% to 1.46% (**Figure 4B**, **Table 4**). For 10 out of the 13 markers, the mill homogenisation step reduced the CV% between replicates, while for heptanoic acid, 4-methyl valeric acid and MPO, the CVs were increased. Mill-homogenising the faeces reduced the CV% for total SCFAs from 20.4% to 7.53%, and for total BCFAs from 15.9% to 7.82%. The CV% for calprotectin decreased from 15.5% to 6.43%, while MPO increased from 24.2% to 48.7%. For none of the markers a statistical difference in the mean concentration values between hammered-only samples and mill-homogenised samples was observed (P-values>0.05; **Table 4**).

**Figure 4.**
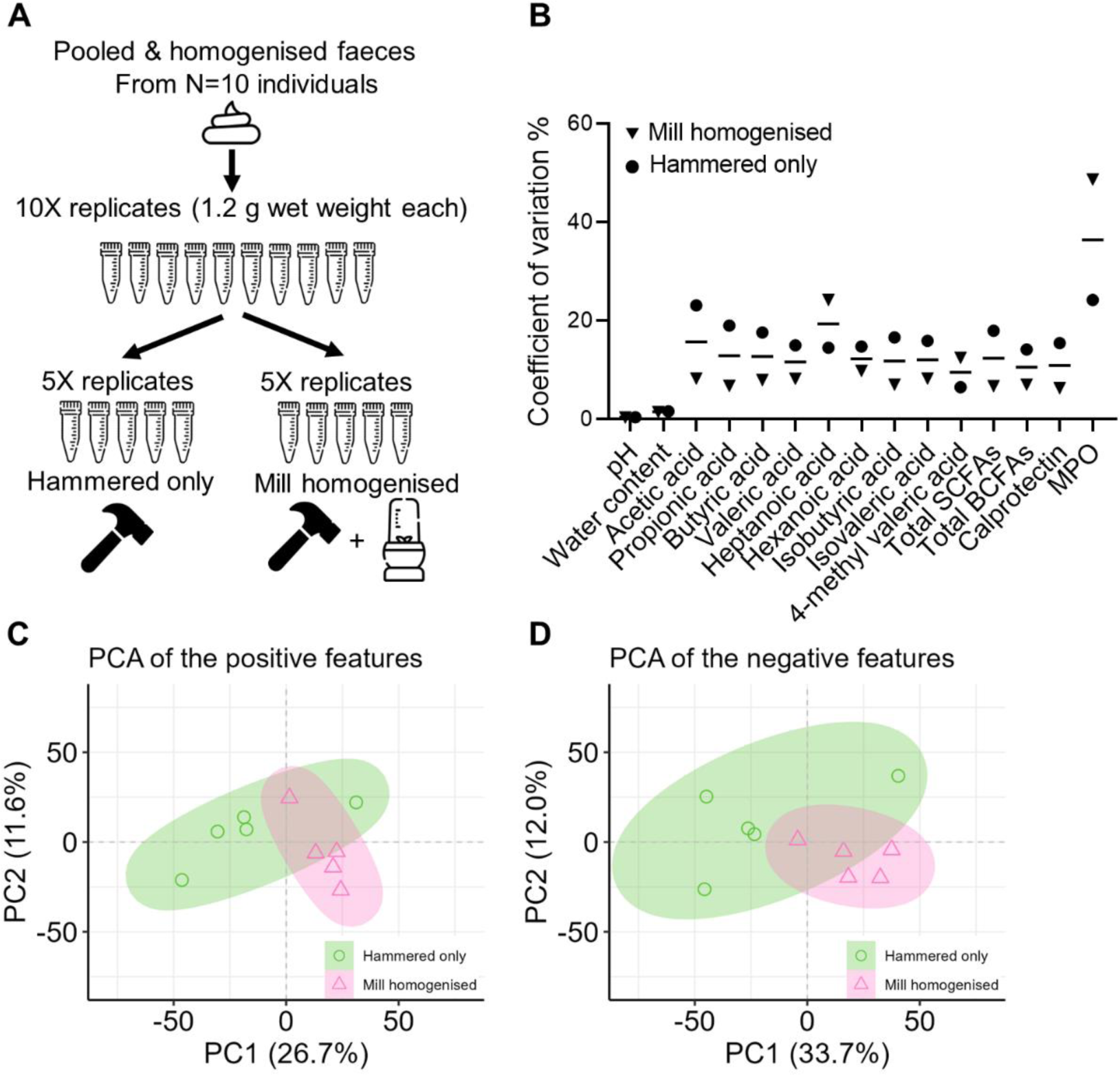
Comparison of variation of faecal pre-processing methods within replicate of the same pooled faecal sample. A) Replicates (n=10) of a pooled faecal sample, of which 5 were pre-processed by hammering-only, and 5 by both hammering and mill-homogenising. B) The coefficient of variation percentage as calculated within the replicates of hammered-only samples (circles) and within replicates of mill-homogenised samples (triangles). C, D) The untargeted metabolomics PCA scores plot based on features detected in positive ionization mode (C) and in negative ionization mode (D) of the hammered-only (circles) and mill-homogenised replicates (triangles).

**Table 4.**
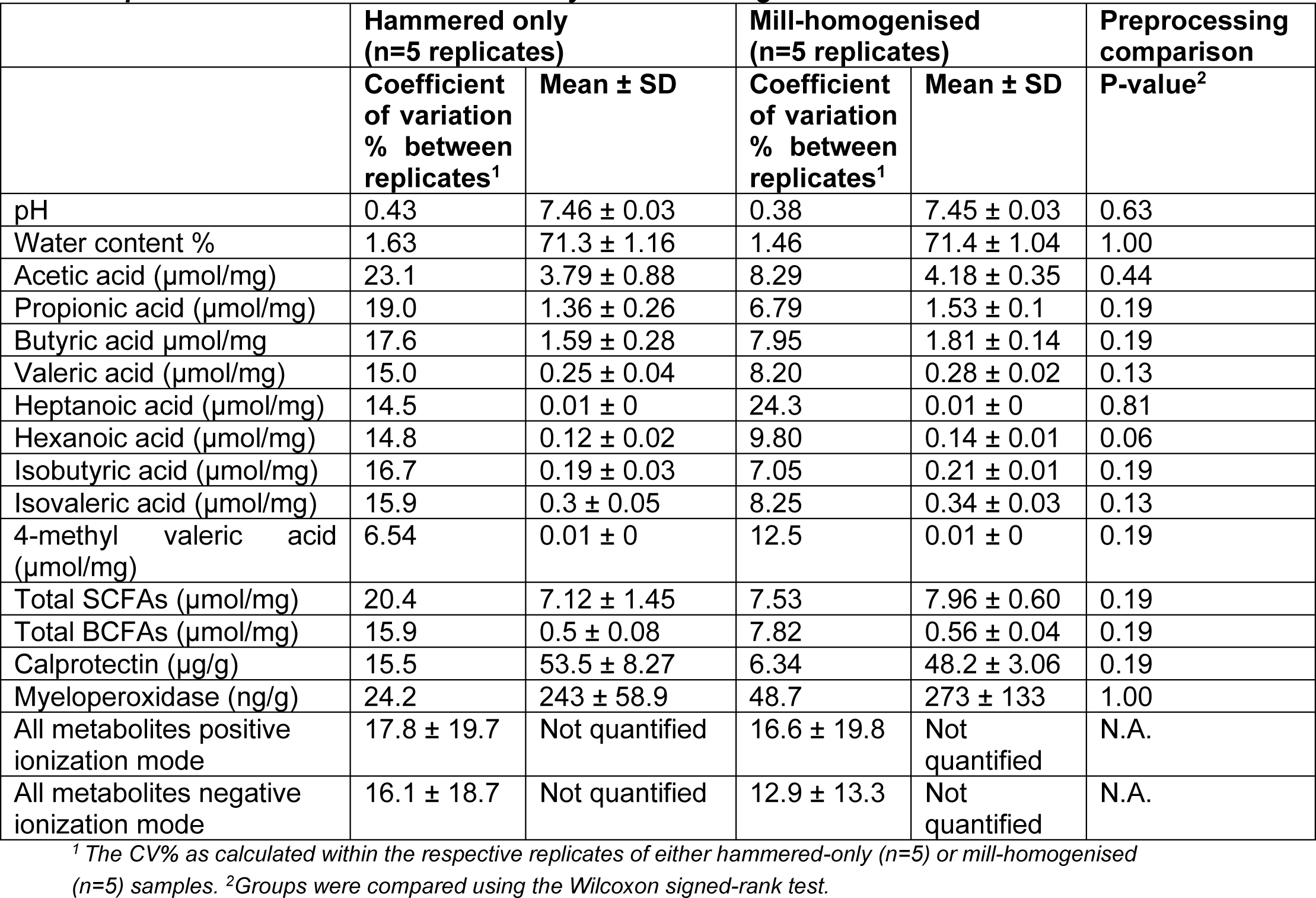
Comparison of faecal pre-processing methods within replicates from a pooled sample that were either hammered-only or mill-homogenised.

|  | Hammered only<br>(n=5 replicates) |  | Mill-homogenised<br>(n=5 replicates) |  | Preprocessing<br>comparison |
| --- | --- | --- | --- | --- | --- |
| | Coefficient<br>of variation<br>% between<br>replicates<sup>1</sup> | Mean ± SD | Coefficient<br>of variation<br>% between<br>replicates<sup>1</sup> | Mean ± SD | P-value<sup>2</sup> |
| pH | 0.43 | 7.46 ± 0.03 | 0.38 | 7.45 ± 0.03 | 0.63 |
| Water content % | 1.63 | 71.3 ± 1.16 | 1.46 | 71.4 ± 1.04 | 1.00 |
| Acetic acid (μmol/mg) | 23.1 | 3.79 ± 0.88 | 8.29 | 4.18 ± 0.35 | 0.44 |
| Propionic acid (μmol/mg) | 19.0 | 1.36 ± 0.26 | 6.79 | 1.53 ± 0.1 | 0.19 |
| Butyric acid μmol/mg | 17.6 | 1.59 ± 0.28 | 7.95 | 1.81 ± 0.14 | 0.19 |
| Valeric acid (μmol/mg) | 15.0 | 0.25 ± 0.04 | 8.20 | 0.28 ± 0.02 | 0.13 |
| Heptanoic acid (μmol/mg) | 14.5 | 0.01 ± 0 | 24.3 | 0.01 ± 0 | 0.81 |
| Hexanoic acid (μmol/mg) | 14.8 | 0.12 ± 0.02 | 9.80 | 0.14 ± 0.01 | 0.06 |
| Isobutyric acid (μmol/mg) | 16.7 | 0.19 ± 0.03 | 7.05 | 0.21 ± 0.01 | 0.19 |
| Isovaleric acid (μmol/mg) | 15.9 | 0.3 ± 0.05 | 8.25 | 0.34 ± 0.03 | 0.13 |
| 4-methyl valeric acid<br>(μmol/mg) | 6.54 | 0.01 ± 0 | 12.5 | 0.01 ± 0 | 0.19 |
| Total SCFAs (μmol/mg) | 20.4 | 7.12 ± 1.45 | 7.53 | 7.96 ± 0.60 | 0.19 |
| Total BCFAs (μmol/mg) | 15.9 | 0.5 ± 0.08 | 7.82 | 0.56 ± 0.04 | 0.19 |
| Calprotectin (μg/g) | 15.5 | 53.5 ± 8.27 | 6.34 | 48.2 ± 3.06 | 0.19 |
| Myeloperoxidase (ng/g) | 24.2 | 243 ± 58.9 | 48.7 | 273 ± 133 | 1.00 |
| All metabolites positive<br>ionization mode | 17.8 ± 19.7 | Not quantified | 16.6 ± 19.8 | Not<br>quantified | N.A. |
| All metabolites negative<br>ionization mode | 16.1 ± 18.7 | Not quantified | 12.9 ± 13.3 | Not<br>quantified | N.A. |

<sup>1</sup> The CV% as calculated within the respective replicates of either hammered-only (n=5) or mill-homogenised (n=5) samples. <sup>2</sup> Groups were compared using the Wilcoxon signed-rank test.

We also investigated the effect of faecal sample homogenisation on untargeted metabolomics and the microbiota composition. The variation in metabolites in the faecal replicates after mill-homogenising was smaller compared to the variation in faecal replicates after hammering only, for both the features detected in positive ionization mode (**Figure 4C**) and the negative ionization mode (**Figure 4D**). For the positive mode, the CV%_intra_ (mean ± SD) of all detected features within the faecal mill-homogenised replicates was 15.0 ± 18.0%, while the hammered only the replicates showed a CV%_intra_ of 16.5 ± 18.3%. For the negative mode features, the CV%_intra_ (mean ± SD) for all metabolites was 12.1 ± 13.1% within the faecal mill-homogenised replicates, while the hammered only replicates had a CV%_intra_ of 15.3 ± 19.3%. To determine the effect of milling on microbiota analysis, first we evaluated the potential fragmentation of DNA, and based on agarose gel analyses it was observed that faecal sample mill-homogenising did not cause DNA fragmentation. The mean (±SD) correlation of the microbiota composition in the hammered only and mill-homogenised faeces at timepoint one within individual was 0.92±0.07 (**Supplementary Table 6**). In comparison, the technical replicates, which was the same microbial DNA included in all consecutive PCR- and sequencing steps, showed a comparable correlation of 0.96±0.03. The overall microbiota composition between the mill-homogenised and hammered-only faeces was not significantly different (PERMANOVA P-value = 0.96), and visualisation of the overall microbiota community did not reveal clusters based on mill-homogenised or hammered-only faeces (**Supplementary Figure 3**). Overall, we showed the added value of the mill-homogenising step for targeted metabolite analyses, since it reduced the variation of SCFAs and BCFAs and untargeted metabolomics in samples obtained from the same faecal pool, and the mill-homogenising did not impact the microbial DNA and composition.

## Discussion

### Variability in gut health markers is marker specific

The main aim of this study was to evaluate the intra-subject variation of selected faecal gut health markers in three faecal samples collected on consecutive days in healthy adults. This marker panel included BSS, pH, WC%, inflammatory biomarkers, total bacteria and fungi copies, microbiota composition and diversity, SCFAs, BCFAs, and untargeted metabolomics. Most markers had a CV%_intra_ less than 30%. This included technical intra-assay variation, which ranged from 0.49-14%. We also reported ICC, but emphasized CV% for interpretation. ICC as a marker of test-retest reliability compares intra-individual variability to inter-individual variation of the sample population, providing a ratio of the variance components. This indicates the relative contributions of different sources of variability. Several markers showed a relatively high intra-individual variability, but nevertheless a good test-retest reliability, due to the ratio with inter-individual variation.

We found that stool consistency, as measured by BSS, had a low intra-individual variability of 16.6% (range 0-43%), comparable to previous findings of 0-57.8% over 9 days in healthy adults [41]. However, our moderate test-retest reliability score (ICC 0.74) contrasts with a study by Vork et al., showing lower test-retest reliability in healthy adults over 7 consecutive days (ICC 0.28) [49]. The BSS test-retest reliability may decrease at the boundaries of constipation (type 2/3) or diarrhoea (type 5/6) [23, 69], so the median stool form type 4 may have contributed to better test-retest reliability in our study. Faecal water content, a more objective measure of stool consistency, had a lower intra-individual variability of 6% (range 1-11%), contrasting with a slightly wider range of 2 – 24% CV%_intra_ in 61 healthy individuals [41]. Despite moderate test-retest reliability (ICC 0.62) in previous literature over 7 consecutive days in healthy controls [49], we found poor test-retest reliability (ICC 0.37) likely due to the relatively low inter-individual variation in comparison to the intra-individual variability. Overall, we confirmed low variation when using BSS scores in healthy participants with mostly stool consistency types 3 or 4. Additionally, a single measure of water content is sufficient, and offers less variability than BSS. Studies that wish to account for stool consistency may consider including faecal water content as objective and less variable measure. Furthermore, faecal pH, reflecting distal colonic pH [39], showed a low intra-individual variability (1.3-6.7%), comparable to findings elsewhere (0.3-8.1%), showing that faecal pH was the most stable over 9 days compared to other gut markers [41]. We also found a that intra-individual variability in pH was positively correlated with intra-individual variability in some faecal SCFAs. We conclude that a single faecal pH measurement is sufficient due to minimal variation and overall stability.

There was moderate intra-individual variability in the concentrations of faecal acetate, propionate, and butyrate. Of these three SCFAs, butyrate showed the highest intra-individual variability (27.8%), followed by propionic acid (17.8%), and acetic acid (16.0%). These findings are consistent with previous findings in faeces collected over three days [70]. Heptanoic, hexanoic, and isovaleric acid had intra-individual variabilities above 30%. For heptanoic acid, the higher variability may be explained by the very low faecal concentrations. Others reported that faecal SCFAs and BCFAs concentrations fluctuated considerably from day-to-day (CV_Intra_ range 26-40%), with valerate varying the least and acetate the most [41]. We found a significant positive correlation between intra-individual variability of faecal water content and intra-individual variation of total SCFAs and butyric acid, highlighting the relevance of including faecal water content as a potential confounding factor in metabolite measurements, also demonstrated by others [71, 72]. BCFAs showed more variable intra-individual responses than SCFAs (CV%_intra_ of 27.4% versus 17%). A single faecal measurement likely suffices for acetate, propionate, butyrate, or total SCFAs and BCFAs, with water content correction potentially reducing butyrate variability.

Furthermore, we found high intra-individual (67%) and inter-individual (222%) variation in total fungi copies. Analysis of intra-individual variability in fungi is sparse, however, previous studies indicate that a large number of the faecal fungi is transient [73–75], mainly derived from dietary consumption and passage from the oral cavity and skin [73, 74, 76–78]. While we did not investigate variability of specific fungi species, others indicated that the occurrence of the same fungi in faeces of healthy volunteers at two timepoints 13-16 weeks apart was less than 20% [73]. We similarly demonstrated high intra-individual variability of 40% (range 13-59%) in total bacteria copies, but with notably lower inter-individual variation (55%) compared to total fungi copies. Our results are similar to findings by Procházková et al, demonstrating an intra-individual variability range of 7.6-72.7 % in total faecal bacterial load over 9 consecutive days, despite their use of a different quantification technique, namely flow cytometry [41]. The high variability may be explained by intestinal transit time that strongly correlates with bacterial mass [79, 80], but we could not confirm a correlation between variability of total bacteria and stool consistency. Additionally, correction for faecal dry weight compared to wet weight did not affect total bacteria and fungi counts variability. Overall, caution is warranted in interpreting single measurements of total bacteria and fungi abundances, and averaging total bacteria and fungi copies from multiple samples is recommended. Further investigation into specific bacteria and fungi abundances within- and between individuals is warranted.

Both inflammatory markers, calprotectin (63.8%) and MPO (106.5%), showed high intra-individual variability. Calprotectin’s variability aligns with previous studies in inflammatory bowel disease (IBD) patients (variabilities of 54% within four days, 39% within two days, and 36% within two days [81–83]) and healthy controls (38% in two non-consecutive samples 1-2 weeks apart [45]). Due to demonstration of diurnal variation by calprotectin, some studies recommend always sampling from the first stool sample of the day to reduce variation [84]. We did not account for the time of day in faecal sampling. However, stool frequency in our study was typically no more than one sample per day, and other studies have shown reliability of calprotectin concentrations regardless of sampling in the morning or evening [82]. Other factors that may affect calprotectin concentrations variability are stool volume and hydration [81], and stool consistency [84]. In our study, we found a significant correlation between the variability of calprotectin and faecal water content, which suggests the importance of including stool consistency in explaining the day-to-day calprotectin variation, also according to international consensus [85]. We conclude that more than one sample of calprotectin is likely needed to accurately quantify calprotectin levels. Further investigation into correcting calprotectin concentrations by faecal water content to account for intra-individual variability is needed. The high intra-individual variability of MPO was consistent with previous variability (20-80%), likely due to difficulties with extracting tightly-bound MPO from the solid stool fraction [86]. In our healthy participants we expected low levels of intestinal inflammation, and indeed only one individual had detectable MPO concentrations in all faecal samples, while five participants had only detectible levels at two timepoints. Therefore, the reliability of the variability of MPO in healthy individuals might not be well illustrated in our small sample size. Despite this, our results indicate that gut inflammation as measured by MPO, is likely to be highly variable in individuals. Future studies incorporating MPO may require multiple faecal samples for accurate interpretation of gut inflammation.

Next, we analysed the variability in the microbiota composition and diversity, and in the overall metabolome using untargeted LCMS-based metabolomics. We demonstrated an overall mean CV%_intra_ of 57% for all microbiota genera within one week. Previously, substantial intra-individual ecological variability longitudinally over 6 weeks to 6 years was seen [48, 87, 88]. In line with our results, most genera showed significant day-to-day variability within individuals, with 72% of all genera demonstrating more than a tenfold change in abundance between consecutive faecal samples [48]. We similarly demonstrated that intra-individual variability is genus-specific. Notably, genera with the highest relative abundance, *Prevotella, Alistipes, Roseburia, Akkermansia, Parabacteroides, Bifidobacterium, Dialister, Clostridium sensus stricto 1, Holdemanella, UCG-002, Sutterrella, Banesiella* demonstrated high intra-individual variability of above 30%. *Clostridium_sensu_stricto_1 (Clostridium ss1)* [89–91] demonstrated the highest intra-individual variability (84%), followed by *Akkermansia* (82%). As yet, there are no publications describing the variation of *Clostridium ss1* as a cluster. Recent interest in the probiotic potential of *Clostridium butyricum,* one species of this cluster, may however, lead to further investigation [92, 93]. *Akkermansia* was the only genus in the top 25 genera with a low test-retest reliability measure (ICC 0.48). *A. muciniphilia* is frequently cited due to its association with beneficial health outcomes [94, 95]. Despite the composition variability, in our study the overall microbial diversity remained stable, which is in line with previous findings demonstrating the stability of intra-individual alpha-diversity over one week [49], but in contrast to substantial alpha-diversity Shannon index changes in individuals over six weeks (33%) [48]. We conclude that despite the relative stability in microbial community, large genus-specific variation is seen. In comparison to the high intra-individual microbiota composition variation, the untargeted metabolomics results showed lower variation, with mean CV%_intra_ of 40% and 41% for features detected in positive and negative ionization mode, respectively. Less than half of all features had CV%_intra_of <30%, indicating a relatively high day-to-day variation in faecal metabolite composition within individuals. The intrinsic dynamics of the microbiota, as well as external factors such as habitual diet, could introduce bias in clinical studies [48, 87, 88]. We demonstrated that this bias may be pronounced on a day-to-day basis when considering specific genera individually. Studies targeting *Akkermansia* as well as other genera with high intra-individual variability, such as *Bifidobacterium,* which are of interest in pre- and probiotic studies, should consider multiple timepoint sampling for accurate assessment. Additionally, future research should confirm intra-individual variability in absolute abundances of these genera, as it may differ from relative abundances [48].

Overall, the intra-individual variability of most of the targeted gut-health markers within subjects was lower than the inter-individual variation between individuals, indicating that the inferences made from analysis of a single timepoint faecal sample should be reliable in the general population. Additionally, we propose that when considering single sampling versus multiple sampling at specific timepoints in studies, the specific gut-health outcomes of primary interest in a study should be taken into account. If genera or markers with high intra-individual coefficient of variation are the primary focus of a study, averaging multiple samples may provide a more accurate assessment of intervention effects, compared to relying on a single faecal sample only.

### Mill-homogenisation during faecal pre-processing decreased variability of gut health markers in faecal replicates

We assessed the impact of an improved sample pre-processing procedure, including additional homogenisation of the frozen hammered faeces into a fine powder using a dedicated liquid nitrogen-cooled mill, on the outcome of measured gut health markers. Compared to sample hammering-only, the additional mill-homogenisation step reduced the CV% between replicates from the same faecal pool for most targeted gut health markers, except for heptanoic acid, 4-methyl valeric acid, and MPO. Our findings corroborate previous studies indicating that homogenisation can decrease the variability in microbiota composition [68, 70], and SCFA levels [70]. Importantly, there were no significant differences in mean marker concentrations between replicates processed by hammering-only and those subjected to mill-homogenisation.

Previous methods, such as manual homogenisation with sterile plastic scoops [70], hand homogenisation [72], homogenisation in saline buffers using commercial grinders [96], or preparing faecal slurries [97] are likely less labour-intensive than our proposed mill-homogenisation method. However, in contrast to our method, which includes a mill that can be cooled with liquid-nitrogen, these other methods do not allow homogenisation of deep-frozen faeces and thus intrinsically incorporate potentially destructive freeze-thaw cycles. Additionally, another study found reduced variability by homogenising entire stool samples in liquid nitrogen and subsampling from the crushed powder, also allowing them to be kept frozen [68]. Hsieh et al. compared randomly sampled fresh and frozen stools that were either not milled at all or milled in a blender or homogenised with a pneumatic mixer, and concluded that homogenisation by any method reduced the microbiota variability [64]. So, for a reduction in measurement variability an adequate sample homogenisation step is preferred and both fresh and frozen stools can be used. However, to prevent microbiome and metabolome changes after sampling, stools should be frozen as soon as possible and then always kept frozen.

Storage of homogenised fine faecal powders, rather than as complete stools or roughly hammered faecal pieces, allows multiple representative aliquoting of the same sample for complementary and repeated analyses. Avoiding thawing of stool samples before microbe or metabolite extraction is key to prevent any thawing-induced metabolic activity and related changes in the microbiome and metabolome of the specific sample. For microbiome analysis using DNA, the currently used grinder (an IKA A11 Basic Analytical Mill) required thorough cleaning between faecal samples, which is time consuming. Alternative grinders like the IKA Tube Mill 100 control in combination with disposable plastic grinding chambers pre-filled with dry ice could streamline this process. Regarding the two protein markers, mill-homogenisation had no significant impact on their concentrations, while the CV% was decreased for calprotectin and increased for MPO, as compared to hammering-only.

Previously, for other tissue matrixes (carcinoma tissues) ball milling was compared to classic tissue pulverization using mortar and pestle, and no differences in protein quality and concentrations were observed [98]. The differential effects of mill-homogenisation on the CVs for protein inflammation markers need further validation and verification by also analysing other proteins that are relevant for gut health, such as immunoglobulin A, lipocalin, and β-defensins [99]. Previously, it has been shown that the intra-stool CV% of calprotectin in three single stool punches ranged between 13-27% (median 17%), of which the authors concluded that a single-stool punch would be reliable for faecal calprotectin measurement [83]. We found a similar CV% for this protein for the hammered-only samples, but showed that mill-homogenising has the potential to further reduce the CV% to only 6%. Overall, mill-homogenisation added value, particularly in the analysis of faecal metabolites, by reducing variation of SCFAs, BCFAs, and untargeted metabolites within samples obtained from the same faecal pool. The proposed milling method can eliminate potential subsampling bias arising from heterogeneous distribution of some gut health markers in faeces. This finding also underscores the importance of applying adequate sample pre-processing techniques to avoid potential erroneous conclusions regarding study outcomes.

## Limitations of the study

Study strengths included the comprehensive assessment of a series of gut health markers as well as both the microbiota and the global metabolome of human stools, which is particularly innovative within the context of intra-individual variability studies. More in-depth insights in the intra-individual stool variability is also key to studies focusing on inter-individual variability. A diverse range of gut health markers, both established and less frequently measured like total bacteria and fungi, provided a multifaceted understanding of gut health marker variability. The optimised faecal sampling procedure, from collection to pre-processing into frozen powders, ensured that the observed variability can be primarily attributed to individual differences rather than methodological and analytical inconsistencies. Furthermore, the proposed mill-homogenisation method was validated using pooled faecal samples, and offers standardization benefits for future research. However, limitations of the present study include a relatively small participant group with mainly female participants, thereby limiting the generalizability and highlighting the necessity for future studies to validate currently obtained results in larger gender-balanced cohorts. Another limitation is that the exact time of faecal collection was not recorded, making it impossible to quantify the time interval between faecal sample collection and diurnal variation. Additionally, in healthy adults with minimal gut inflammation, markers such as calprotectin and MPO may have limited relevance. The current focus on sampling stools on consecutive days overlooks potential within-day variations, as shown previously by others [70], limiting our understanding of even shorter-term variability. The lack of detailed background information on the volunteers, including dietary records and medication use, does not allow for explanation of potential causes of the observed variation in assessed gut health markers. Known factors influencing variability of microbial metabolites and other health markers include for instance metabolite absorption rates in the intestine, GI transit time, responses to diet, microbiota composition, colonic pH, and time since last meal [37–39, 73, 80, 100, 101]. In the present study we only included water content and stool consistency as proxies for transit time, where future studies should include direct measures [102] to account for microbiota and metabolite variation. Future research aiming for a more comprehensive analysis should aim to incorporate these factors potentially influencing gut marker variation.

## Conclusions

In conclusion, our study revealed that most of the selected gut-health markers in stools exhibited a low intra-individual variability, except for total fungi, total bacteria, calprotectin, MPO and certain microbiota genera. Notably, these included the more commonly measured genera such as *Prevotella_9*, *Bifidobacterium*, *Roseburia* and *Akkermansia.* Microbiota alpha-diversity within individual remained stable over a week of consecutive stool sampling, and microbiota variability was genus-dependent. Our findings help determine whether a single faecal sample or multiple samples are needed for accurate marker representation. Additionally, our optimised sample pre-processing, including mill-homogenisation of frozen stools, effectively reduced the variation in faecal metabolites. Therefore, we conclude that faecal mill-homogenisation mitigates the analytical variability in metabolites and eliminates subsampling bias that may result from a heterogeneous distribution of certain gut health markers in human faecal samples.

## Methodology

### Study participants and ethics approval

Ten healthy adults were recruited from Wageningen, the Netherlands. Exclusion criteria were antibiotic use within the three months before study start, and the presence of gastrointestinal disorders. All participants gave written informed consent. The Medical Ethical Reviewing Committee of Wageningen University (METC-WU) evaluated this study and concluded that this research does not fall within the remit of the Dutch Medical Research Involving Human Subjects act.

### Faecal sample collection

Participants collected faecal samples on three consecutive days, or alternatively at three timepoints as close as possible to each other within one week. The collection occasions can be found in **Supplementary Table 1**. Participants collected faeces at home using a faecal collection paper (Faecesvanger, Tag Hemi VOF, The Netherlands) and 55×44 mm polypropylene collection containers with a collection scoop integrated into the screw cap (Sarstedt, Nümbrecht, Germany). Participants were instructed to collect from three different topographical locations of the faeces, and to fill the container until two-thirds. Directly after collection, samples were stored at -20°C at home, for maximally seven days. Afterwards, samples were transported under frozen conditions, in Styrofoam containers with ice packs, to the research facility and were immediately stored at -80°C upon arrival. Participants indicated for each faecal sample the date and timepoint of sample collection, the number of stools passed per day, and the stool consistency on a Bristol Stool Scale (BSS). This scale describes seven types of stools ranging from 1: hard/lumpy to 7: watery without solid pieces.

### Faecal sample processing

Faecal samples were processed under frozen conditions in two phases. First, frozen samples were crushed with a dead-blow hammer (Performance Tool Dead-Blow Hammer, Wilmar Corporation) into smaller particles of approximately 0.5 cm^2^.During the hammering, samples were always kept frozen using dry ice and liquid nitrogen. The hammered sample was stored at -80°C, and 20 grams of the hammered faeces was additionally mill-homogenised in a IKA A11 Basic Analytical Mill (Sigma-Aldrich, Darmstadt, Germany). Samples were always kept on dry ice. Liquid nitrogen was applied to the samples during both the hammering and milling steps to ensure they remained in a deeply-frozen state. Between homogenisation of samples, the mill was thoroughly cleaned using milli-q water and ethanol. Liquid nitrogen and dry ice were used to ensure the faeces always remained deeply frozen. Processed samples were stored at -80°C. Subsequent aliquoting and weighing faeces for consecutive analyses were also performed under frozen conditions. Samples were thawed only if indicated for specific downstream analysis. For all analyses, except for the protein biomarker analyses, mill-homogenised samples were used. All faecal sample analyses from the same individual for all measurements were performed within the same batch.

### Evaluation of the effect of mill homogenisation using pooled faeces

To determine whether IKA milling affected outcome measures, 10 replicates of 1-2 gram faeces were made from a pooled sample of 15 gram faeces obtained from all ten participants. Afterwards, five replicates were then processed by hammering only, while five replicates were both hammered and mill-homogenised. For all markers of interest, comparison was made between the CV% within the five hammered-only replicates or the five mill-homogenised replicates. For the microbiota comparison only, the mill-homogenised samples and hammered-only samples of timepoint 1 of all participants were compared. To determine the effect of milling on DNA integrity, DNA was isolated from hammered-only and mill-homogenised samples. Afterwards, the DNA was checked on agarose gels (1%) using electrophoresis (100 V, 60 min) and ethidium bromide to check potential DNA degradation.

### Water content and pH measurement

For faecal water content measurement 1 gram of mill-homogenised faecal sample was placed on an aluminium dish (VWR Aluminium smooth weigh dish 57mm, VWR International, USA) and dried at 95°C for a minimum of 48 hours by vacuum drying (Heraeus D-6450 vacuum oven). Samples were weighed before and after drying. The water content percentage was determined as follows: 100-[dry weight (g)/wet weight (g)*100]. For faecal pH measurement 1 gram of mill-homogenised faecal sample was diluted in 10 mL milli-q water. Samples were homogenised for 15 min on a horizontal shaker (Universal shaker SM-30, Edmund Bühler GmbH, Bodelshausen, Germany) and centrifuged (5292 *xg*, 10 min, RT) before measurement. The pH of the supernatant was measured using a PCE-228-R meter (PCE Instruments, Enschede, The Netherlands) modified on the method described elsewhere [103].

### Protein biomarkers

Concentrations of faecal protein biomarkers were measured using enzyme-linked immunosorbent assay (ELISA) for MPO (Immundiagnostik, Bensheim, Germany), and calprotectin (Immundiagnostik) according to manufacturer’s instructions. For each assay, 60±6 mg of hammered-only faeces was dissolved in extraction buffer (IDK Extract, Immundiagnostik) and further processed according to manufacturer’s instructions.

### Short chain and branched chain fatty acids

The SCFAs (acetic acid, propionic acid, butyric acid, valeric acid, heptanoic acid, hexanoic acid) and BCFAs (isobutyric acid, isovaleric acid, 4-methylvaleric acid) levels were measured using gas chromatography with flame ionization detection (GC-FID) using 8690 GC system (Agilent, Amstelveen, the Netherlands) and a Restek Stabilwax-DA column (Restek Corporation, United States) as described by Lotti et al. [104] with minor adjustments. In short, 40-50 mg of mill-homogenised faeces was weighed off in duplicate and 990 μL methyl tert-butyl ether, 10 μL 15% phosphoric acid in milli-q water and 10 μL 2-ethylbutyric acid in MTBE as the internal standard (IS) was added. Samples were homogenised for 30 min at 1300 rpm on an orbital shaker (IKA VXR basic Vibrax®, IKA®-Werke GmbH & Co. KG, Staufen, Germany) and centrifuged (3000xg, 10 min, RT). 200 μL of the organic phase layer was then transferred to an injection vial. Injection volume on the GC-FID was 2 μl with an additional split ratio of 1:10. A final volume of 0.1 μL was injected into the column. The GC run settings were as follows: i). 1 min at 80°C, ii.) raised by 10 °C /min to 200°C, iii.) hold time of 5 minutes iv.) raised by 25 °C/min, to a temperature of 230 °C, v.) 5 minutes hold time. The total run time was 24.2 min. The carrier gas was nitrogen at a flow rate of 1.32ml/min. A 6-point calibration curve was prepared from standards in MTBE: acetic acid (2.47-600 mM), propionic acid (0.49-120 mM), iso-butyric acid (0.25-60 mM), butyric acid (0.99-240 mM), isovaleric acid (0.25-60 mM), valeric acid (0.25-60 mM), 4-methyl valeric acid (isohexanoic/isocaproic acid) (0.25-60 mM), hexanoic acid (0.25-60 mM), heptanoic acid (0.25-60 mM), and IS was set at 40 mM. The fatty acids were identified using a reference standard mix (article CRM46975, Sigma-Aldrich). This reference mix was also used to verify the accuracy of the calibration curve, which showed a deviation of <5% for all compounds. A pooled faecal sample was analysed in duplicate in each analytical batch to monitor the quality of the analyses. The inter-batch CV% values ranged between 6 and 14% for all compounds. Concentrations of the SCFAs were calculated per mg of wet weight.

### Untargeted metabolomics

Approximately 80 mg of freeze-dried mill-homogenised faeces was used for untargeted metabolomics analysis using liquid chromatography–mass spectrometry (LC-MS). UHPLC-Q-Orbitrap-MS/MS analysis was conducted using an Vanquish UHPLC (Thermo Fisher Scientific, Carlsbad, CA, USA) connected to a Q-Exactive Focus™ mass spectrometer (Thermo Scientific, Sunnyvale, CA, USA) with heated electrospray ionization (HESI-II; Thermo Fisher Scientific). The stool extracts were chromatographically separated on an Waters ACQUITY

HSS T3 UPLC column (2.1 × 100 mm, 3.0 μm) using a mobile phase consisting of 0.1% formic acid/water (A) and acetonitrile (B). Gradient elution was performed as: 0–1 min, 5% B; 1–8 min, 5–95% B; 8–11 min, 95% B; 11-12, 95-5% B and 12-15 min, 5% B. The column was maintained at 40 °C, the flow rate was 0.4 mL/min and the injection volume was 2 μL. The mass spectrum data was acquired in polarity switching mode positive and negative ion mode through full MS. Representative samples were also acquired using data-dependent MS/MS. Control software used was Xcalibur, version 4.2.27). The scan parameters were set as follows: scan range, 80 to 1000 m/z; resolution, 70000; AGC target, 1 × 106; maximum inject time, 100 ms. The following parameters were used for the heated electrospray ionization source (HESI source): spray voltage, +/-3.5 kV for ESI− and ESI+ in polarity switching mode; sheath gas flow rate, 40 arb; auxiliary gas flow rate, 10 arb; sweep gas flow rate, 0 arb; capillary temperature, 320 °C.

Six quality controls (QC), which consisted of a pool of the faecal samples from the participants, were analysed to ensure precision and stability of the LC-MS machine during the whole process of injection. QC samples were injected every after 10 sample injections and was used to assess chromatographic drift and instrumental errors. Features were extracted using the raw spectra as import in open-source software [105, 106]. The detailed processing settings can be found in **Supplementary Table 7**. RANSAC (RANdom SAmple Consensus) alignment of features across all samples was performed. Gap filling was performed on the aligned feature list using the peak finder option. The raw data files including the ID, m/z, retention time, and peak area of the positive- and negative polarity mode features were exported for further pre-processing in MetaboAnalyst 5.0 (https://www.metaboanalyst.ca/). The features with more than 20% of missing values were removed. The remaining missing values were imputed with the limit of detection values using 1/5 of the minimum positive value for each variable. Features with a relative standard deviation (RSD = SD/mean) of over 20% in the quality control samples were removed. Additionally, 40% of the features was filtered using IQR filter. The data pre-processing flowchart of positive and negative polarity modes is shown in **Supplementary Figure 2**. Data was normalized using normalization by sum, log transformation with base 10, and auto scaling. Principal Component Analysis (PCA) was performed on the normalized and scaled metabolomics data.

### Total bacteria and total fungi quantification

The total copy number of the 16S rRNA gene, indicative for the total bacteria, and the total copy number of the 18S rRNA gene, indicative for the total fungi, were quantified using digital droplet PCR (ddPCR, QX200™ Droplet Digital™ PCR System, Bio-Rad Laboratories, CA). All materials and apparatus for the assay were obtained from Bio-Rad Laboratories. Before analysis, the DNA stocks were diluted 2000 times for fungi analyses, and 100000 times for bacteria analyses. The primers and probes for the analysis of total bacteria [107] and total fungi [108] were used as described previously [107, 108]. The PCR reaction mix contained 10 μL QX200™ ddPCR™ Supermix for Probes (no UTP), 1 μL forward primer (18 μM), 1 μL reverse primer (18 μM), 1 μL probe (5 μM), 2 μL nuclease free water, and 5 μL diluted DNA. The final volume of 20 μL was transferred to the DG8 cartridge. Droplet Generator Oil (70 μL) for EvaGreen were also added and the cartridge was covered with DG8 gasket, to generate droplets in the droplet generator (Bio-Rad). The generated droplets (40 μL) were transferred to the PCR plate. The plate was sealed at 170°C for 5 sec (PX1 PCR Plate Sealer, Bio-Rad). Then, the PCR reaction took place in the thermos T100 Thermal Cycler. The cycling protocol was as follows: 95°C for 10 min; 50 cycles of 94°C for 30 sec and 58°C for 2.5 min; 98°C for 10 min; and 4°C for 30 min. The plate was placed in the Droplet Reader (Bio-Rad). QuantaSoft Analysis Pro was used for the data analysis.

### Microbiota composition and predicted functionality

Faecal DNA was extracted from 250±20 mg faeces using the DNeasy Powersoil Pro Kit (Qiagen, Venlo, the Netherlands) according to manufacturer’s instructions. The variable V4 region of the 16S rRNA gene was amplified in a PCR reaction using barcoded primers, and afterwards PCR products were purified, quantified and pooled in equimolar concentrations as described in detail elsewhere [109]. The PCR products were subjected to paired-end sequencing using the Illumina Novaseq6000 platform (Novogene, Cambridge, UK). The sequencing data was processed using NG-Tax 2.0 using default settings [110]. Raw sequencing data were demultiplexed by trimming barcodes and primer sequences. The SILVA reference database version 138 was used to assign taxonomy. R version 4.2.1 was used for the microbiota analyses. Mitochondrial filtering was applied. In total, 1391 amplicon sequencing variants (ASVs) were detected, and 216 genera. The median number of reads per sample was 194292 (IQR 152269, 219321). Raw counts were transformed to relative abundance for intra- and inter-individual variability analyses. The alpha-diversity, the within-sample diversity, was calculated on the read counts using the microbiome package using the indexes Inverse Simpson (considers number of species and evenness), Shannon (considers abundance and evenness), and Fisher (associated with the number of species). Also the Faith phylogenetic diversity, which is based on branch length connecting taxa in those samples and the root node of the phylogenetic tree, was calculated using the picante package. Beta-diversity, the between-sample diversity, using proportion normalized abundances of all amplicon sequence variants using weighted UniFrac distance as implemented in the phyloseq package to determine overall microbiota differences, and visualised using Principal Coordinate Analysis (PCoA). Permutational multivariate analysis of variance (PERMANOVA) as implemented in the vegan package was used to determine differences in the overall microbiota community between milled and non-milled faeces. Principal Coordinate Analysis (PCoA) with Weighted UniFrac using the microbiota data on ASV level was performed using the R package *phyloseq*. The microbiota functional pathways related to SCFA production were predicted based on the 16S rRNA gene sequences using the phylogenetic investigation of communities by reconstruction of unobserved states algorithm version 2.5.0 (PICRUSt2) with default settings [111], but with an adjusted minimum alignment of 50%. The predicted pathways and enzymes were transformed to relative abundance. Pathways classes and enzymes related to SCFA production were selected from the MetaCyc database using Pathway Tools version 26.0 [112].

### Statistical analysis

Statistical analyses were performed in R version 4.2.1. Test-retest reliability of markers over the three timepoints were assessed using intra-class correlation coefficients (ICC). The ICC evaluates the variance as the variance within individuals in relation to the total variance, where total variance is the variance within and between individuals. For our analysis, the ICCs were computed using R package *irr* with a two-way mixed effects model using absolute agreement between timepoints based on single measurements [113]; 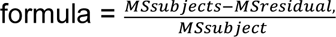 where the MSsubjects indicates the mean squares due to the subjects as is calculated by comparing the variance of subject means to the grand mean, and MSresidual indicates the residual mean square reflecting variance (represents the within-subject variability). ICC values indicated test-retest reliability as follows: < 0.5 poor, 0.5 to 0.75 moderate, 0.75 to 0.9 good, and > 0.90 excellent [113]. Pairwise Pearson’s or Spearman’s correlations, depending on the normality of distribution, were calculated to compare all markers between the three timepoints within individual, between the milled and non-milled faeces, and between technical replicates. Coefficient of variation (CV%, formula: [(*σ / μ) × 100%*]) were calculated between consecutive timepoints within individuals (CV%_intra_) and between individuals (CV%_inter_) per timepoint, and within and between replicates from the pooled faeces, either hammered-only or mill-homogenised, quality controls. A cut-off of 30% was considered as high biological variation. Statistical significance was accepted as P<0.05.

## Acknowledgements

First, we thank all volunteers who donated stool samples for this study. We would like to thank the Gut Health Club colleagues Anna Malinowska and Marie-Luise Puhlmann for the input on the design of the study. Also we thank Soumya Kar for the input on the set-up of the study. We would like to thank Geert Althuizen, Yueyi Zhong, and Panos Fourmouzis for assisting with the laboratory and data analysis. We would also like to thank Maria-João Paulo for her advice on statistical analysis. But above all, we would like to thank our beloved mentor Wilma Steegenga, who passed away before this manuscript was finalised. She initiated and set-up this study, which she was so passionate about, and she inspired us all with her expertise, scientific rigor, and ambition. She is immensely missed.

This project was partially funded by the Horizon 2020 Framework Programme of the European Union (ITN SmartAge, H2020-MSCA-ITN-2019-859890). The contents of this publication are the sole responsibility of the authors and do not necessarily reflect the opinion of the European Union. K.K. and Y.M were funded by the Horizon 2020 Framework Programme of the European Union (ITN SmartAge, H2020-MSCA-ITN-2019-859890). M.v.T. was funded by a Crossover grant (MOCIA 17611) of the Dutch Research Council (NWO). The MOCIA programme is a public-private partnership (see https://mocia.nl/scientific/).

## Author contributions

Conceptualization, W.S, D.K, G.J.H, R.d.V, G.B.G. and M.v.T.; Methodology and Investigation, K.K., Y.M., N.v.d.W., S.K., R.d.V, M.G.B., K.M., M.D.d.B, M.B., N.L., G.B.G, W.S., and M.v.T.; Funding Acquisition, L.d.G. and W.S.; Supervision, L.d.G. and Y.V.; Writing original draft, K.K., M.v.T., and W.S.; Writing-Review and editing - All authors. All authors, except W.S. who passed away before finalization of the manuscript, approved the final version.

## Ethics declaration

The Medical Ethical Reviewing Committee of Wageningen University (METC-WU) evaluated this study and concluded that this research does not fall within the remit of the Dutch Medical Research Involving Human Subjects Act. The subjects gave written consent for the use of their faeces. All specimens were used and coded anonymously.

## References

1. Ahlawat, S., Asha, and K.K. Sharma, Gut-organ axis: a microbial outreach and networking. Lett Appl Microbiol, 2021. 72(6): p. 636–668.

2. Lama Tamang, R., et al., The diet-microbiota axis: a key regulator of intestinal permeability in human health and disease. Tissue Barriers, 2022: p. 2077069.

3. Ghosh, T.S., et al., Adjusting for age improves identification of gut microbiome alterations in multiple diseases. Elife, 2020. 9.

4. Durack, J. and S.V. Lynch, The gut microbiome: Relationships with disease and opportunities for therapy. J Exp Med, 2019. 216(1): p. 20–40.

5. He, Y., et al., Regional variation limits applications of healthy gut microbiome reference ranges and disease models. Nat Med, 2018. 24(10): p. 1532–1535.

6. Yatsunenko, T., et al., Human gut microbiome viewed across age and geography. Nature, 2012. 486(7402): p. 222–7.

7. Asnicar, F., et al., Microbiome connections with host metabolism and habitual diet from 1,098 deeply phenotyped individuals. Nat Med, 2021. 27(2): p. 321–332.

8. Campaniello, D., et al., How Diet and Physical Activity Modulate Gut Microbiota: Evidence, and Perspectives. Nutrients, 2022. 14(12).

9. Huang, X., et al., Dietary variety relates to gut microbiota diversity and abundance in humans. Eur J Nutr, 2022.

10. Aya, V., et al., Association between physical activity and changes in intestinal microbiota composition: A systematic review. PLoS One, 2021. 16(2): p. e0247039.

11. Dziewiecka, H., et al., Physical activity induced alterations of gut microbiota in humans: a systematic review. BMC Sports Sci Med Rehabil, 2022. 14(1): p. 122.

12. Vich Vila, A., et al., Impact of commonly used drugs on the composition and metabolic function of the gut microbiota. Nat Commun, 2020. 11(1): p. 362.

13. Walsh, J., et al., Drug-gut microbiota interactions: implications for neuropharmacology. Br J Pharmacol, 2018. 175(24): p. 4415–4429.

14. Wilmanski, T., et al., Gut microbiome pattern reflects healthy ageing and predicts survival in humans. Nat Metab, 2021. 3(2): p. 274–286.

15. Wu, Y.L., et al., Gut microbiota alterations and health status in aging adults: From correlation to causation. Aging Med (Milton), 2021. 4(3): p. 206–213.

16. Manor, O., et al., Health and disease markers correlate with gut microbiome composition across thousands of people. Nat Commun, 2020. 11(1): p. 5206.

17. Procházková, N., et al., Advancing human gut microbiota research by considering gut transit time. Gut, 2022.

18. Vandeputte, D., et al., Stool consistency is strongly associated with gut microbiota richness and composition, enterotypes and bacterial growth rates. Gut, 2016. 65(1): p. 57–62.

19. Tropini, C., How the Physical Environment Shapes the Microbiota. mSystems, 2021. 6(4): p. e0067521.

20. Zhernakova, A., et al., Population-based metagenomics analysis reveals markers for gut microbiome composition and diversity. Science, 2016. 352(6285): p. 565–9.

21. Hammer, J. and S.F. Phillips, Fluid loading of the human colon: effects on segmental transit and stool composition. Gastroenterology, 1993. 105(4): p. 988–98.

22. Lewis, S.J. and K.W. Heaton, Stool form scale as a useful guide to intestinal transit time. Scand J Gastroenterol, 1997. 32(9): p. 920–4.

23. Blake, M.R., J.M. Raker, and K. Whelan, Validity and reliability of the Bristol Stool Form Scale in healthy adults and patients with diarrhoea-predominant irritable bowel syndrome. Aliment Pharmacol Ther, 2016. 44(7): p. 693–703.

24. Nordin, E., et al., Modest Conformity Between Self-Reporting of Bristol Stool Form and Fecal Consistency Measured by Stool Water Content in Irritable Bowel Syndrome and a FODMAP and Gluten Trial. Am J Gastroenterol, 2022. 117(10): p. 1668–1674.

25. Galazzo, G., et al., How to Count Our Microbes? The Effect of Different Quantitative Microbiome Profiling Approaches. Front Cell Infect Microbiol, 2020. 10: p. 403.

26. Knight, R., et al., Best practices for analysing microbiomes. Nat Rev Microbiol, 2018. 16(7): p. 410–422.

27. Vandeputte, D., et al., Quantitative microbiome profiling links gut community variation to microbial load. Nature, 2017. 551(7681): p. 507–511.

28. Gutierrez, M.W. and M.C. Arrieta, The intestinal mycobiome as a determinant of host immune and metabolic health. Curr Opin Microbiol, 2021. 62: p. 8–13.

29. Shuai, M., et al., Mapping the human gut mycobiome in middle-aged and elderly adults: multiomics insights and implications for host metabolic health. Gut, 2022.

30. Zhang, L., et al., The role of gut mycobiome in health and diseases. Therap Adv Gastroenterol, 2021. 14: p. 17562848211047130.

31. Chen, H., et al., A Forward Chemical Genetic Screen Reveals Gut Microbiota Metabolites That Modulate Host Physiology. Cell, 2019. 177(5): p. 1217–1231.e18.

32. Krautkramer, K.A., J. Fan, and F. Bäckhed, Gut microbial metabolites as multi-kingdom intermediates. Nat Rev Microbiol, 2021. 19(2): p. 77–94.

33. Schroeder, B.O. and F. Bäckhed, Signals from the gut microbiota to distant organs in physiology and disease. Nat Med, 2016. 22(10): p. 1079–1089.

34. Visconti, A., et al., Interplay between the human gut microbiome and host metabolism. Nat Commun, 2019. 10(1): p. 4505.

35. Zheng, X., X. Cai, and H. Hao, Emerging targetome and signalome landscape of gut microbial metabolites. Cell Metab, 2022. 34(1): p. 35–58.

36. Blaak, E., et al., Short chain fatty acids in human gut and metabolic health. Beneficial microbes, 2020. 11(5): p. 411–455.

37. Rios-Covian, D., et al., An Overview on Fecal Branched Short-Chain Fatty Acids Along Human Life and as Related With Body Mass Index: Associated Dietary and Anthropometric Factors. Front Microbiol, 2020. 11: p. 973.

38. Ramos Meyers, G., H. Samouda, and T. Bohn, Short Chain Fatty Acid Metabolism in Relation to Gut Microbiota and Genetic Variability. Nutrients, 2022. 14(24).

39. Yamamura, R., et al., Intestinal and fecal pH in human health. Frontiers in Microbiomes, 2023. 2.

40. Firrman, J., et al., The impact of environmental pH on the gut microbiota community structure and short chain fatty acid production. FEMS Microbiology Ecology, 2022. 98(5).

41. Procházková, N., et al., Gut environmental factors explain variations in the gut microbiome composition and metabolism within and between healthy adults. bioRxiv, 2024: p. 2024.01.23.574598.

42. LaBouyer, M., et al., Higher total faecal short-chain fatty acid concentrations correlate with increasing proportions of butyrate and decreasing proportions of branched-chain fatty acids across multiple human studies. Gut Microbiome, 2022. 3.

43. Prata, M.d.M.G., et al., Comparisons between myeloperoxidase, lactoferrin, calprotectin and lipocalin-2, as fecal biomarkers of intestinal inflammation in malnourished children. Journal of translational science, 2016. 2(2): p. 134.

44. Gubatan, J., et al., Antimicrobial peptides and the gut microbiome in inflammatory bowel disease. World Journal of Gastroenterology, 2021. 27(43): p. 7402.

45. Padoan, A., et al., Improving IBD diagnosis and monitoring by understanding preanalytical, analytical and biological fecal calprotectin variability. Clin Chem Lab Med, 2018. 56(11): p. 1926–1935.

46. Hansberry, D.R., et al., Fecal Myeloperoxidase as a Biomarker for Inflammatory Bowel Disease. Cureus, 2017. 9(1): p. e1004.

47. Johnson, A.J., et al., Daily Sampling Reveals Personalized Diet-Microbiome Associations in Humans. Cell Host & Microbe, 2019. 25(6): p. 789–802.e5.

48. Vandeputte, D., et al., Temporal variability in quantitative human gut microbiome profiles and implications for clinical research. Nat Commun, 2021. 12(1): p. 6740.

49. Vork, L., et al., Does Day-to-Day Variability in Stool Consistency Link to the Fecal Microbiota Composition? Front Cell Infect Microbiol, 2021. 11: p. 639667.

50. David, L.A., et al., Host lifestyle affects human microbiota on daily timescales. Genome Biol, 2014. 15(7): p. R89.

51. Turroni, S., et al., Temporal dynamics of the gut microbiota in people sharing a confined environment, a 520-day ground-based space simulation, MARS500. Microbiome, 2017. 5(1): p. 39.

52. Venkataraman, A., et al., Variable responses of human microbiomes to dietary supplementation with resistant starch. Microbiome, 2016. 4(1): p. 33.

53. McOrist, A.L., et al., Bacterial population dynamics and faecal short-chain fatty acid (SCFA) concentrations in healthy humans. British Journal of Nutrition, 2008. 100(1): p. 138–146.

54. Karu, N., et al., A review on human fecal metabolomics: Methods, applications and the human fecal metabolome database. Anal Chim Acta, 2018. 1030: p. 1–24.

55. Hosseinkhani, F., et al., Towards Standards for Human Fecal Sample Preparation in Targeted and Untargeted LC-HRMS Studies. Metabolites, 2021. 11(6).

56. Johnson, A.J., et al., A Guide to Diet-Microbiome Study Design. Front Nutr, 2020. 7: p. 79.

57. De Saedeleer, B., et al., Systematic characterization of human gut microbiome-secreted molecules by integrated multi-omics. ISME Commun, 2021. 1: p. 82.

58. Deda, O., et al., Sample preparation optimization in fecal metabolic profiling. J Chromatogr B Analyt Technol Biomed Life Sci, 2017. 1047: p. 115–123.

59. El Manouni El Hassani, S., et al., Optimized sample preparation for fecal volatile organic compound analysis by gas chromatography-mass spectrometry. Metabolomics, 2020. 16(10): p. 112.

60. Gratton, J., et al., Optimized Sample Handling Strategy for Metabolic Profiling of Human Feces. Anal Chem, 2016. 88(9): p. 4661–8.

61. Lamichhane, S., et al., Gut metabolome meets microbiome: A methodological perspective to understand the relationship between host and microbe. Methods, 2018. 149: p. 3–12.

62. Liang, Y., et al., Systematic Analysis of Impact of Sampling Regions and Storage Methods on Fecal Gut Microbiome and Metabolome Profiles. mSphere, 2020. 5(1).

63. Trošt, K., et al., Describing the fecal metabolome in cryogenically collected samples from healthy participants. Sci Rep, 2020. 10(1): p. 885.

64. Hsieh, Y.H., et al., Impact of Different Fecal Processing Methods on Assessments of Bacterial Diversity in the Human Intestine. Front Microbiol, 2016. 7: p. 1643.

65. Jones, J., et al., Fecal sample collection methods and time of day impact microbiome composition and short chain fatty acid concentrations. Sci Rep, 2021. 11(1): p. 13964.

66. Wu, W.K., et al., Optimization of fecal sample processing for microbiome study - The journey from bathroom to bench. J Formos Med Assoc, 2019. 118(2): p. 545–555.

67. Bruce, K., et al., A practical guide to DNA-based methods for biodiversity assessment. 2021: Pensoft Advanced Books.

68. Gorzelak, M.A., et al., Methods for Improving Human Gut Microbiome Data by Reducing Variability through Sample Processing and Storage of Stool. PLOS ONE, 2015. 10(8): p. e0134802.

69. Chumpitazi, B.P., et al., Bristol Stool Form Scale reliability and agreement decreases when determining Rome III stool form designations. Neurogastroenterol Motil, 2016. 28(3): p. 443–8.

70. Jones, J., et al., Fecal sample collection methods and time of day impact microbiome composition and short chain fatty acid concentrations. Scientific Reports, 2021. 11(1): p. 13964.

71. Siigur, U., et al., Concentrations and Correlations of Faecal Short-chain Fatty Acids and Faecal Water Content in Man. Microbial Ecology in Health and Disease, 1994. 7(6): p. 287–294.

72. McOrist, A.L., et al., Bacterial population dynamics and faecal short-chain fatty acid (SCFA) concentrations in healthy humans. British journal of nutrition, 2008. 100(1): p. 138–146.

73. Hallen-Adams, H.E., Fungi inhabiting the healthy human gastrointestinal tract: a diverse and dynamic community. Fungal Ecology, 2015. 15: p. 9–17.

74. Hallen-Adams, H.E. and M.J. Suhr, Fungi in the healthy human gastrointestinal tract. Virulence, 2017. 8(3): p. 352–358.

75. Raimondi, S., et al., Longitudinal Survey of Fungi in the Human Gut: ITS Profiling, Phenotyping, and Colonization. Front Microbiol, 2019. 10: p. 1575.

76. Auchtung, T.A., et al., Investigating Colonization of the Healthy Adult Gastrointestinal Tract by Fungi. mSphere, 2018. 3(2): p. 10.1128/msphere.00092-18.

77. Underhill, D.M. and I.D. Iliev, The mycobiota: interactions between commensal fungi and the host immune system. Nat Rev Immunol, 2014. 14(6): p. 405–16.

78. David, L.A., et al., Diet rapidly and reproducibly alters the human gut microbiome. Nature, 2014. 505(7484): p. 559–63.

79. Stephen, A.M., H.S. Wiggins, and J.H. Cummings, Effect of changing transit time on colonic microbial metabolism in man. Gut, 1987. 28(5): p. 601–9.

80. Daniel, H., Diet and Gut Microbiome and the “Chicken or Egg” Problem. Frontiers in Nutrition, 2022. 8.

81. Tibble, J., et al., A simple method for assessing intestinal inflammation in Crohn’s disease. Gut, 2000. 47(4): p. 506–513.

82. Kristensen, V., et al., Clinical importance of faecal calprotectin variability in inflammatory bowel disease: intra-individual variability and standardisation of sampling procedure. Scandinavian Journal of Gastroenterology, 2016. 51(5): p. 548–555.

83. Cremer, A., et al., Variability of Faecal Calprotectin in Inflammatory Bowel Disease Patients: An Observational Case-control Study. Journal of Crohn’s and Colitis, 2019. 13(11): p. 1372–1379.

84. Lasson, A., et al., The Intra-Individual Variability of Faecal Calprotectin: A Prospective Study In Patients With Active Ulcerative Colitis. Journal of Crohn’s and Colitis, 2014. 9(1): p. 26–32.

85. D’Amico, F., et al., International consensus on methodological issues in standardization of fecal calprotectin measurement in inflammatory bowel diseases. United European Gastroenterol J, 2021. 9(4): p. 451–460.

86. Swaminathan, A., et al., Faecal Myeloperoxidase as a Biomarker of Endoscopic Activity in Inflammatory Bowel Disease. Journal of Crohn’s and Colitis, 2022. 16(12): p. 1862–1873.

87. Olsson, L.M., et al., Dynamics of the normal gut microbiota: A longitudinal one-year population study in Sweden. Cell Host Microbe, 2022. 30(5): p. 726–739.e3.

88. Zhou, X., et al., Longitudinal profiling of the microbiome at four body sites reveals core stability and individualized dynamics during health and disease. Cell Host & Microbe, 2024. 32(4): p. 506–526.e9.

89. Gupta, R.S. and B. Gao, Phylogenomic analyses of clostridia and identification of novel protein signatures that are specific to the genus Clostridiumsensu stricto (cluster I). International Journal of Systematic and Evolutionary Microbiology, 2009. 59(2): p. 285–294.

90. Li, C.-J., et al., Comparative genomic analysis and proposal of Clostridium yunnanense sp. nov., Clostridium rhizosphaerae sp. nov., and Clostridium paridis sp. nov., three novel Clostridium sensu stricto endophytes with diverse capabilities of acetic acid and ethanol production. Anaerobe, 2023. 79: p. 102686.

91. Wiegel, J., Tanner, R., Rainey, F.A, The Prokaryotes Volume 4. Bacteria: Firmicutes, Cyanobacteria. An Introduction to the family Clostridiaceae., ed. M. Dworkin. Vol. 4. New York: Springer.

92. Liu, L., et al., Clostridium butyricum Potentially Improves Immunity and Nutrition through Alteration of the Microbiota and Metabolism of Elderly People with Malnutrition in Long-Term Care. Nutrients, 2022. 14(17).

93. Sun, Y.Y., et al., The effect of Clostridium butyricum on symptoms and fecal microbiota in diarrhea-dominant irritable bowel syndrome: a randomized, double-blind, placebo-controlled trial. Sci Rep, 2018. 8(1): p. 2964.

94. Geerlings, S.Y., et al., Akkermansia muciniphila in the Human Gastrointestinal Tract: When, Where, and How? Microorganisms, 2018. 6(3).

95. Zhang, T., et al., Akkermansia muciniphila is a promising probiotic. Microbial Biotechnology, 2019. 12(6): p. 1109–1125.

96. Ezzy, A.C., et al., Storage and handling of human faecal samples affect the gut microbiome composition: A feasibility study. Journal of Microbiological Methods, 2019. 164: p. 105668.

97. Gangadoo, S., et al., The Multiomics Analyses of Fecal Matrix and Its Significance to Coeliac Disease Gut Profiling. Int J Mol Sci, 2021. 22(4).

98. Ostapowicz, J., et al., Comparison of tumour tissue homogenisation methods: mortar and pestle versus ball mill. 2023.

99. Siddiqui, I., H. Majid, and S. Abid, Update on clinical and research application of fecal biomarkers for gastrointestinal diseases. World J Gastrointest Pharmacol Ther, 2017. 8(1): p. 39–46.

100. McOrist, A.L., et al., Fecal butyrate levels vary widely among individuals but are usually increased by a diet high in resistant starch. J Nutr, 2011. 141(5): p. 883–9.

101. Roager, H.M. and L.O. Dragsted, Diet-derived microbial metabolites in health and disease. Nutrition Bulletin, 2019. 44(3): p. 216–227.

102. Asnicar, F., et al., Blue poo: impact of gut transit time on the gut microbiome using a novel marker. Gut, 2021. 70(9): p. 1665–1674.

103. Million, M., et al., Increased Gut Redox and Depletion of Anaerobic and Methanogenic Prokaryotes in Severe Acute Malnutrition. Sci Rep, 2016. 6: p. 26051.

104. Lotti, C., et al., Development of a fast and cost-effective gas chromatography–mass spectrometry method for the quantification of short-chain and medium-chain fatty acids in human biofluids. Analytical and Bioanalytical Chemistry, 2017. 409(23): p. 5555–5567.

105. Tsugawa, H., et al., A lipidome atlas in MS-DIAL 4. Nature Biotechnology, 2020. 38(10): p. 1159–1163.

106. Schmid, R., et al., Integrative analysis of multimodal mass spectrometry data in MZmine 3. Nature Biotechnology, 2023. 41(4): p. 447–449.

107. Liu, C.M., et al., BactQuant: An enhanced broad-coverage bacterial quantitative real-time PCR assay. BMC Microbiology, 2012. 12(1): p. 56.

108. Liu, C.M., et al., FungiQuant: A broad-coverage fungal quantitative real-time PCR assay. BMC Microbiology, 2012. 12(1): p. 255.

109. Rios-Morales, M., et al., A toolbox for the comprehensive analysis of small volume human intestinal samples that can be used with gastrointestinal sampling capsules. Scientific Reports, 2021. 11(1): p. 1–14.

110. Poncheewin, W., et al., NG-Tax 2.0: a semantic framework for high-throughput amplicon analysis. Frontiers in Genetics, 2020. 10: p. 1366.

111. Langille, M.G., et al., Predictive functional profiling of microbial communities using 16S rRNA marker gene sequences. Nat Biotechnol, 2013. 31(9): p. 814–21.

112. Caspi, R., et al., The MetaCyc database of metabolic pathways and enzymes. Nucleic Acids Research, 2017. 46(D1): p. D633–D639.

113. Koo, T.K. and M.Y. Li, A Guideline of Selecting and Reporting Intraclass Correlation Coefficients for Reliability Research. J Chiropr Med, 2016. 15(2): p. 155–63.

