## Supplementary Information for "An optimised approach to evaluate variability in gut health markers in healthy adults"

***Supplementary Table 1. Faecal sampling frequency and stool consistency of the ten healthy subjects.***

| Participant | Timepoint | Number of stools per day | Collected stools | BSS |
|---|---|---|---|---|
| 1 | 1 | 1 | Day 1 | 3 |
| | 2 | 1 | Day 2 | 2 |
| | 3 | 1 | Day 3 | 2 |
| 2 | 1 | 1 | Day 1 | 4 |
| | 2 | 2 | Day 5 | 4 |
| | 3 | 3 | Day 8 | 4 |
| 3 | 1 | Not stated | Day 1 | 5 |
| | 2 | Not stated | Day 2 | 6[1] |
| | 3 | Not stated | Day 3 | 6 |
| 4 | 1 | 1 | Day 1 | 5[1] |
| | 2 | Not stated | Day 2 | 6[1] |
| | 3 | 1 | Day 3 | 4 |
| 5 | 1 | 2 | Day 1 | 4 |
| | 2 | 2 | Day 2 | 4 |
| | 3 | 2 | Day 3 | 4 |
| 6 | 1 | 1 | Day 1 | 2 |
| | 2 | 1 | Day 2 | 2 |
| | 3 | 1 | Day 4 | 4 |
| 7 | 1 | 1 | Day 1 | 5[1] |
| | 2 | 1 | Day 2 | 3[1] |
| | 3 | 1 | Day 3 | 3 |
| 8 | 1 | 2 | Day 1 | 6 |
| | 2 | 2 | Day 3 | 5 |
| | 3 | 2 | Day 4 | 6 |
| 9 | 1 | 1 | Day 1 | 2[1] |
| | 2 | 1 | Day 2 | 3 |
| | 3 | 1 | Day 3 | 2 |
| 10 | 1 | 1 | Day 1 | 3[1] |
| | 2 | 3 | Day 2 | 3 |
| | 3 | 2 | Day 3 | 3 |

*[1]Where participants chose two BSS scores, the highest of the two were included in the analysis.*
*BSS: Bristol Stool Scale.*

***Supplementary Table 2. The intra-class correlation coefficients of the predicted pathways and enzymes related to SCFA production in faecal samples of healthy subjects collected over three consecutive days[1].*** *The microbiota functionality was derived from the total predicted genomes, predicted based on the 16S rRNA gene sequencing data.*

| | | **Coefficient of variation % [2]** | | |
|---|---|---|---|---|
| **Microbial predicted gene or pathway** | **EC number or pathway number** | **Intra-individual ($CV\%_{intra}$)** | **Inter-individual ($CV\%_{inter}$)** | **Intra-class correlation coefficient [CI] [1]** |
| **Acetate production** | | | | |
| *Acetate kinase* | EC 2.7.2.1 | 2.93 ± 3.01 | 12.2 ± 2.04 | 0.91 [0.77-0.97] |
| *Phosphate acetyltransferase* | EC 2.3.1.8 | 2.05 ± 2.20 | 5.76 ± 0.79 | 0.75 [0.46-0.92] |
| *Acetate ligase* | EC 6.2.1.1 | 18.7 ± 15.1 | 44.6 ± 6.86 | 0.75 [0.46-0.92] |
| | EC 6.2.1.13 | 74.8 ± 42.0 | 246.1 ± 60.8 | 0.66 [0.31-0.89] |
| *Pyruvate fermentation to acetate and lactate II* | PWY-5100 | 4.44 ± 3.87 | 9.90 ± 1.68 | 0.71 [0.39-0.91] |
| *Acetylene degradation (anaerobic)* | P161-PWY | 8.23 ± 4.56 | 19.2 ± 2.10 | 0.82 [0.57-0.95] |
| *Bifidobacterium shunt* | P124-PWY | 22.2 ± 16.2 | 50.2 ± 2.16 | 0.73 [0.42-0.91] |
| *Hexitol fermentation to lactate, formate, ethanol and acetate* | P461-PWY | 17.9 ± 7.92 | 67.2 ± 2.69 | 0.94 [0.83-0.98] |
| *L-glutamate degradation V (via hydroxyglutarate)* | P162-PWY | 39.1 ± 30.3 | 96.1 ± 10.9 | 0.85 [0.64-0.96] |
| *L-lysine fermentation to acetate and butanoate* | P163-PWY | 33.4 ± 26.1 | 58.2 ± 7.66 | 0.72 [0.41-0.91] |
| **Propionate production** | | | | |
| *Propionyl-coA synthetase* | EC 6.2.1.17 | 46.5 ± 34.8 | 91.4 ± 25.0 | 0.39 [0.03-0.76] |
| *Propionate coA-transferase* | EC 2.8.3.1 | 15.8 ± 9.36 | 61.9 ± 3.71 | 0.88 [0.71-0.97] |
| *Propionaldehyde dehydrogenase* | EC 1.2.1.87 | 17.2 ± 14.26 | 81.2 ± 6.86 | 0.89 [0.71-0.97] |
| *Pyruvate fermentation to propanoate I* | P108-PWY | 14.4 ± 9.25 | 38.2 ± 1.84 | 0.81 [0.57-0.94] |
| *(S)-propane-1,2-diol degradation* | PWY-7013 | 23.2 ± 16.1 | 42.3 ± 7.69 | 0.71 [0.37-0.91] |
| **Butyrate production** | | | | |
| *Butyrate kinase* | EC 2.7.2.7 | 8.16 ± 5.47 | 16.3 ± 1.60 | 0.68 [0.35-0.90] |
| *Acetate CoA-transferase α subunit* | EC 2.8.3.8 | 19.9 ± 9.55 | 34.7 ± 7.91 | 0.63 [0.26-0.88] |
| *Acetate CoA-transferase β subunit* | EC 2.8.3.9 | 20.8 ± 8.16 | 35.6 ± 8.94 | 0.63 [0.26-0.88] |
| *4-aminobutanoate degradation V* | PWY-5022 | 22.0 ± 14.7 | 81.7 ± 3.64 | 0.86 [0.67-0.96] |

| | | | | |
|---|---|---|---|---|
| *Acetyl-CoA fermentation to butanoate* | PWY-5676 | 14.1 ± 6.42 | 29.55 ± 8.68 | 0.74 [0.43-0.92] |
| *L-glutamate degradation V (via hydroxyglutarate)* | P162-PWY | 39.1 ± 30.3 | 96.12 ± 10.90 | 0.85 [0.64-0.96] |
| *L-lysine fermentation to acetate and butanoate* | P163-PWY | 33.4 ± 26.1 | 58.2 ± 7.66 | 0.72 [0.41-0.91] |
| *Pyruvate fermentation to butyrate* | CENTFERM-PWY | 28.5 ± 20.9 | 44.7 ± 12.8 | 0.45 [0.08-0.79] |
| *Succinate fermentation to butanoate* | PWY-5677 | 22.2 ± 7.40 | 52.8 ± 10.5 | 0.83 [0.58-0.95] |
| **Lactate production** | | | | |
| *L-lactate dehydrogenase* | EC 1.1.1.27 | 9.62 ± 6.47 | 21.1 ± 2.45 | 0.76 [0.48-0.93] |
| *D-lactate dehydrogenase* | EC 1.1.1.28 | 4.93 ± 2.31 | 28.5 ± 2.62 | 0.96 [0.89-0.99] |

*[1] The 95% confidence interval is provided. [2] The mean ±SD of the n=10 participants is shown. [3] Spearman's correlations between the three timepoint were analysed pairwise and the correlation range is shown.*

***Supplementary Table 3. Variability of total fungi and bacteria counts corrected for faecal wet weight or dry weight in faecal samples of healthy subjects collected over three consecutive days.***

| **Variable** | **Coefficient of variation % [1]** | | **Intra-class correlation coefficient [CI][2]** |
|---|---|---|---|
| | **Intra-individual ($CV\%_{intra}$)** | **Inter-individual ($CV\%_{inter}$)** | |
| Total fungi copies corrected for mg faecal wet weight | 67.0±35.8 | 222.0±16.7 | 0.90 [0.75-0.97] |
| Total fungi copies corrected for mg faecal dry weight | 64.1±39.0 | 225.6±24.6 | 0.95 [0.86-0.99] |
| Total bacteria copies corrected for mg faecal wet weight | 40.6±16.9 | 55.0±18.9 | 0.32 [-0.01-0.71] |
| Total bacteria copies corrected for mg faecal dry weight | 41.1±24.4 | 58.0±21.9 | 0.32 [-0.01-0.71] |

***Supplementary Table 4. Technical intra- and inter-assay variation of the measured gut health markers.***

| **Measurement** | **Technical variation (coefficient of variation %)** | |
|---|---|---|
| | **Intra-assay CV%** | **Inter-assay CV%** |
| BSS | NA | NA |
| pH | 0.53 | NA |
| Water content | 0.49 | 0.17 |
| Acetic acid | 3.07 | 8.08 |
| Propionic acid | 3.31 | 7.55 |
| Butyric acid | 3.15 | 7.16 |
| Heptanoic acid | 5.13 | 14.03 |
| Hexanoic acid | 3.54 | 8.24 |
| Valeric acid | 2.83 | 6.91 |
| Isobutyric acid | 3.28 | 7.44 |
| Isovaleric acid | 3.19 | 7.49 |
| 4-methyl valeric acid | 5.49 | 11.50 |
| Total fungi | ND | ND |
| Total bacteria | ND | ND |
| Calprotectin | 13.99 | 9.80[1] |
| Myeloperoxidase | 13.67 | 10.95[1] |

*[1]As only one assay was run, the mean inter-assay CV% as per technical specifications of commercial assay kit is provided. NA, not applicable. ND, not determined.*

***Supplementary Table 5. The variability of all microbiota measured on genus level as measured with the coefficient of variation and the inter-class correlation coefficient.***

| **Intra-individual [2] variability (n=number of bacteria on genus level detected)** | **Mean CV%$_{intra}$ of detected bacteria ± SD** | **Mean CV%$_{intra}$ of detected bacteria ± SD (*n*=10 individuals)** | **% of bacteria with <30 CV%** | **Mean % of bacteria with <30 CV% ± SD (*n*=10 individuals)** |
|---|---|---|---|---|
| Subject 1 (n=51) [1] | 51.7±48.0 | 56.5±9.0 | 43.1 | 43.1±14.7 |
| Subject 2 (n=58) | 68.6±45.4 | | 17.2 | |
| Subject 3 (n=53) | 51.7±54.1 | | 50.9 | |
| Subject 4 (n=83) | 49.7±51.0 | | 49.4 | |
| Subject 5 (n=51) | 61.3±45.0 | | 27.5 | |
| Subject 6 (n=75) | 62.5±55.1 | | 40.0 | |
| Subject 7 (n=76) | 67.7±58.8 | | 30.3 | |
| Subject 8 (n=62) | 48.4±43.3 | | 48.4 | |
| Subject 9 (n=55) | 41.8±50.4 | | 65.5 | |
| Subject 10 (n=60) | 61.3±45.0 | | 58.3 | |
| **Intra-class correlation coefficient** | **% of bacteria [1]** | | | |
| Good or excellent test-retest reliability (ICCs 0.75-1.0) | 43.3 | | | |
| Moderate test-retest reliability (ICCs 0.50-0.75) | 29.8 | | | |
| Poor test-retest reliability (ICCs <0.50) | 27.0 | | | |

*[1] The number of bacteria on genus level differed per individual and are mentioned for each subject.*
*[2] Inter-individual variation is not reported, since the variability and the type of bacteria present in the faecal samples between individuals was too high to make meaningful calculations.*

***Supplementary Table 6. The correlation of the microbiota composition in the mill-homogenised and hammered only faecal samples of participants, and of the technical replicates[1].*** *Individual values of the n=10 adults are provided.*

| **Participant** | **Correlation between mill-homogenised and hammered-only faeces of the first timepoint** | **Correlation technical replicates[2]** |
|---|---|---|
| 1 | 0.72 | 0.93 |
| 2 | 0.96 | 0.92 |
| 3 | 0.93 | 0.97 |
| 4 | 0.93 | 0.93 |
| 5 | 0.96 | 0.96 |
| 6 | 0.94 | 0.92 |
| 7 | 0.95 | 0.99 |
| 8 | 0.94 | 0.99 |
| 9 | 0.95 | 0.98 |
| 10 | 0.93 | 0.99 |
| Mean (±SD): | 0.92±0.07 | 0.96±0.03 |

*[1]Spearman correlations were calculated. [2]The technical replicate was the same microbial DNA included in a different PCR reaction.*

***Supplementary Table 7. Summary of the optimised MZmine parameters for processing of the untargeted metabolomics data in the Software MZ-mine 3.***

| **Processing modules** | **Parameters** | **Values** |
|---|---|---|
| Mass detection | Scan types (IMS) | All scan types |
| | Mass detector | Auto |
| Mass detection | Noise level | 2.0E4 |
| Chromatogram building | Minimum group size in number of scans | 5 |
| | Group intensity threshold | 6.0E4 |
| | Minimum highest intensity | 2.0E5 |
| | Scan to scan accuracy | 0.04 m/z or 5 ppm |
| Feature resolving | Chromatographic threshold | 65% |
| | Minimum search range RT/Mobility (absolute) | 0.03 |
| | Minimum relative height | 5% |
| | Minimum absolute height | 2.0E5 |
| | Minimum ratio of peak top/edge | 2 |
| | Peak duration range (min/mobility) | 0-0.5 |
| Isotope filtering | m/z tolerance | 0.02 m/z or 5 ppm |
| | Retention time tolerance | 0.01 absolute (min) |
| | Monotonic shape | √ |
| | Maximum charge | 2 |
| | Representative isotope | Lowest m/z |
| Feature alignment | m/z tolerance | 0.04 m/z or 5ppm |
| | RT tolerance | 0.2 absolute (min) |
| | RT tolerance after correction | 0.05 absolute (min) |
| | RANSAC iterations | 10000 |
| | Minimum number of points | 30% |
| | Threshold value | 0.03 |
| | Linear model | √ |
| Gap-filling | Intensity tolerance | 20% |
| | m/z tolerance | 0.04 m/z or 5 ppm |
| | Retention time tolerance | 0.05 absolute (min) |

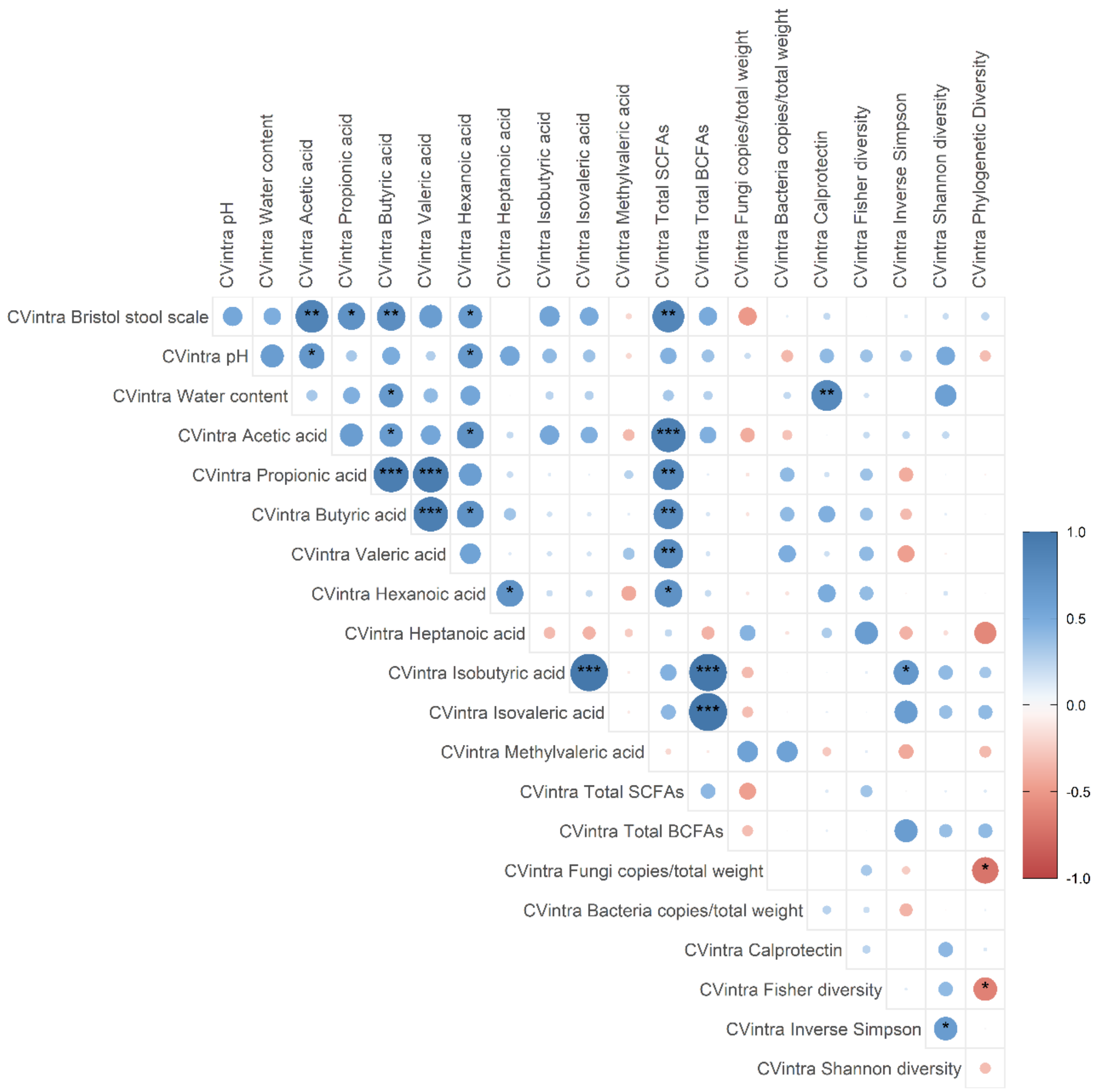


***Supplementary Figure 1. Heatmap of the correlations between the intra-individual variabilities of the selected gut health markers.*** *In total n=10 individuals (intra-CV of the three consecutive faecal samples) were included in the correlation analysis. The scale indicates the Spearman correlation coefficient, ranging from blue r=1.0 to r=-1.0. Significances are shown by *P-values≤0.05, **P-values≤0.01, ***P-values≤0.001 after FDR correction.*

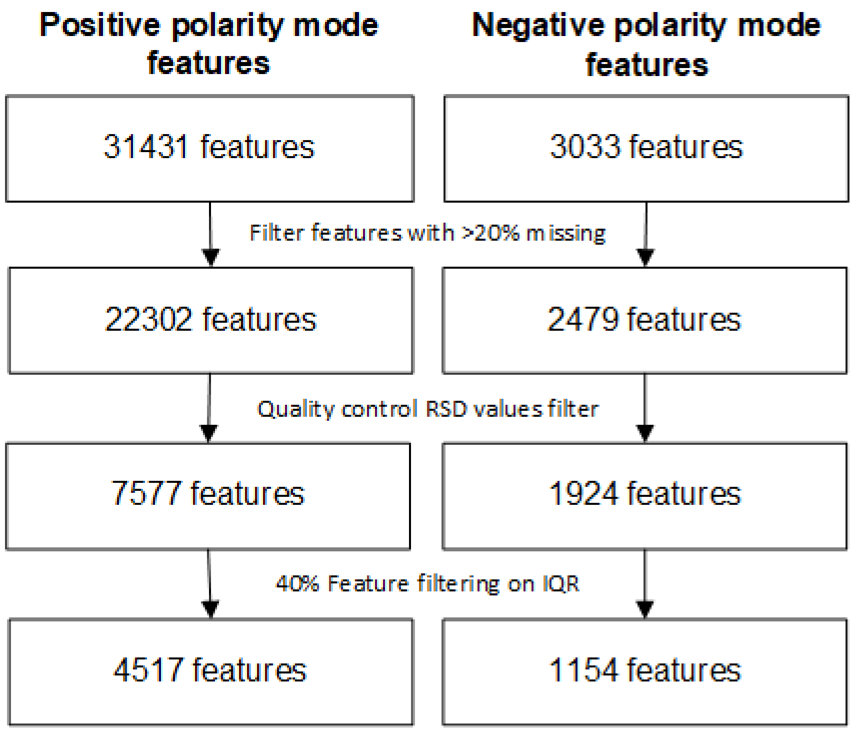


***Supplementary Figure 2. Metabolomics data pre-processing flowchart of positive and negative polarity modes.***

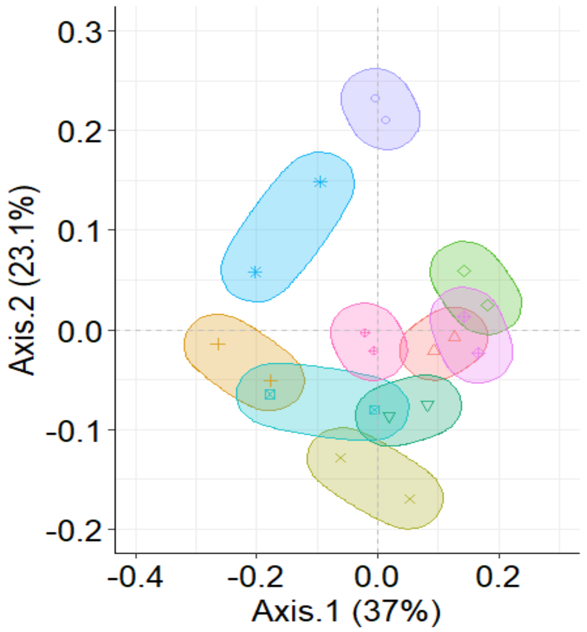


***Supplementary Figure 3. The overall faecal microbiota differences in adults.*** *The beta-diversity was calculated using Weighted UniFrac. The PCoA plot visualises the microbiota variation between the individuals, with one mill-homogenised and hammered-only faecal sample from the same individual. The 95% confidence ellipses are shown. PCoA: Principal Coordinate Analysis.*
